# Integrating multi-covariate disentanglement with counterfactual analysis on synthetic data enables cell type discovery and counterfactual predictions

**DOI:** 10.1101/2025.06.03.657578

**Authors:** Stathis Megas, Arian Amani, Antony Rose, Olli Dufva, Kian Shamsaie, Hesam Asadollahzadeh, Krzysztof Polanski, Muzlifah Haniffa, Sarah A. Teichmann, Mohammad Lotfollahi

## Abstract

Single-cell gene expression is influenced by diverse covariates such as genomics protocol, tissue origin, donor attributes, and microenvironment, which are challenging to disentangle. We present CellDISECT, a novel method combining disentangled representations and causal inference for multi-batch, multi-covariate single-cell data analysis. CellDISECT employs a mixture of expert variational autoencoders to learn covariate-specific and unsupervised latent spaces, enabling counterfactual predictions and biological discovery. Drawing inspiration from LLM training on synthetic data, CellDISECT generates synthetic counterfactuals during training and their quality is scored in the loss function. This semi-autoencoding of counterfactuals during training increases model performance in counterfactual predictions at test time. Benchmarking across datasets, CellDISECT outperformed existing methods in disentanglement, counterfactual *in-silico* prediction of responses to perturbations, and cell type discovery. CellDISECT predicted responses of cells to changing tissue microenvironments and identified a novel pre-natal megakaryocyte subpopulation with immune characteristics distinct from classical platelet-producing MKs, highlighting its unique capabilities in single-cell analysis to help identify novel subpopulations and reduce concerns of technical effects during integration.

## Introduction

A cell’s gene expression profile is a function of multiple attributes such as tissue of origin, its surrounding microenvironment, biometric factors of the donor, and experimental technical variables, as well as clinical variables such as treatments or infections. Single-cell sequencing has enabled profiling gene expression at single-cell resolution in a high-throughput manner^1,2^, but it still remains challenging to analyze and interpret millions of cells at the same time due to the simultaneous and intertwined effects of the aforementioned attributes on gene expression, which can be hard to disentangle^3^. Such a disentangled representation of single-cell data would be more interpretable and would facilitate understanding of biological mechanisms^4,5^, and if combined with causal modeling, would also allow for better counterfactual predictions^6^. Counterfactual predictions could allow understanding the molecular responses of various cell types and patient groups to environmental challenges such as infections or therapeutic interventions.

Disentanglement methods in representation learning aim to find a latent representation that naturally organizes in semantically distinct components^4,5^. Such representations are at the core of explainable AI since they not only facilitate learning but also offer insights into the inner workings of neural networks, allowing us to understand why models make the predictions they do.

In statistics, causal inference is the field that tries to uncover causal links between variables in the presence of spurious correlations in the data^6^. Uncovering these causal links can then be leveraged to predict counterfactual outcomes, which are simulated outcomes for different scenarios and actions that were never observed. Since at any given time, we can only opt for one course of action, often referred to as the fundamental problem of causal inference, counterfactual simulation analysis is necessary to estimate individualized treatment effects and personalize medical treatments^7^.

Finally, recent work in large language modelling has shown opportunities and limitations^8,9^ of training on data generated by AI, often referred to as synthetic data. In the seminal Deepseek-R1 series of models^10^, the authors used Deepseek-R1 to generate 800k (synthetic) training samples to then train a set of smaller models. However, training recursively on synthetically (non-causally) generated data leads to mode collapse^11,12^.

In single-cell data, there have been earlier attempts at disentangled representations with Biolord^13^ and scDisInFact^14^ and also earlier attempts with causal inference methods^15–17^ for single-cell data, but not both simultaneously. Importantly, only one of these methods^16^ leverages the abilities of generative models to be trained on their own (synthetic) data, but it does not perform an ablation study of its effect. Moreover, most of these methods allow modeling of only one covariate at a time^15,16^; learn cell embeddings that are akin to covariate embeddings and are therefore not able to capture biological information^13,14,16^; and suffer from lack of causal identifiability which can impact the quality of the counterfactual predictions^13,14^. Modelling multiple covariates is essential in complex biological datasets for understanding the impact of features such as age, sex and genetics simultaneously on cell states.

Here we introduce Cell DISentangled Experts for Covariate counTerfactuals (CellDISECT) for disentangled representations and causal inference for multi-batch, multi-covariate single-cell data analysis. We demonstrate the utility of CellDISECT in counterfactual predictions of responses to inflammatory stimuli, infectious agents, and tissue microenvironments across cell types and patient groups. We also reprocess and integrate a public single-cell cross-organ study of the developing immune system^18^ to resolve the different populations of immune cell types and revealed a novel subpopulation of early megakaryocytes (MKs) with immune properties but distinct from previously identified immune MKs in the yolk sac (YS)^19^. Moreover, CellDISECT highlighted important differences in erythropoiesis between liver and bone marrow.

## Results

### CellDISECT offers covariate-supervised cell embeddings that capture biological information

Existing approaches for disentangled representations in single cell data are focused on either counterfactual predictions^13^ or inferring which genes are affected by each covariate^14^. As such, they utilize latent spaces resembling covariate embeddings, which cannot be effectively leveraged to explore cell type specificity.

We present CellDISECT, a method that integrates disentangled representations with causal learning on synthetic training data to analyse multi-batch, multi-covariate single-cell datasets. (see **Figure 1a**). CellDISECT consists of a mixture of expert VAEs^20^, each specializing in one of the covariates at a time by finding a representation of cells that is supervised by this covariate as well as an unsupervised VAE that aims to learn a representation for unobserved covariates (see **Figure 1b**). CellDISECT can then leverage the learnt latent factors to compute counterfactual gene expression due to perturbations on one covariate at a time or many covariates together, a novelty that has hitherto been ignored in the literature (see **Figure 1c**). During the training process we repeatedly generate (synthetic) counterfactuals and include a penalty term in the loss function for the quality of these predictions (see Methods).

**Legend Figure 1:**
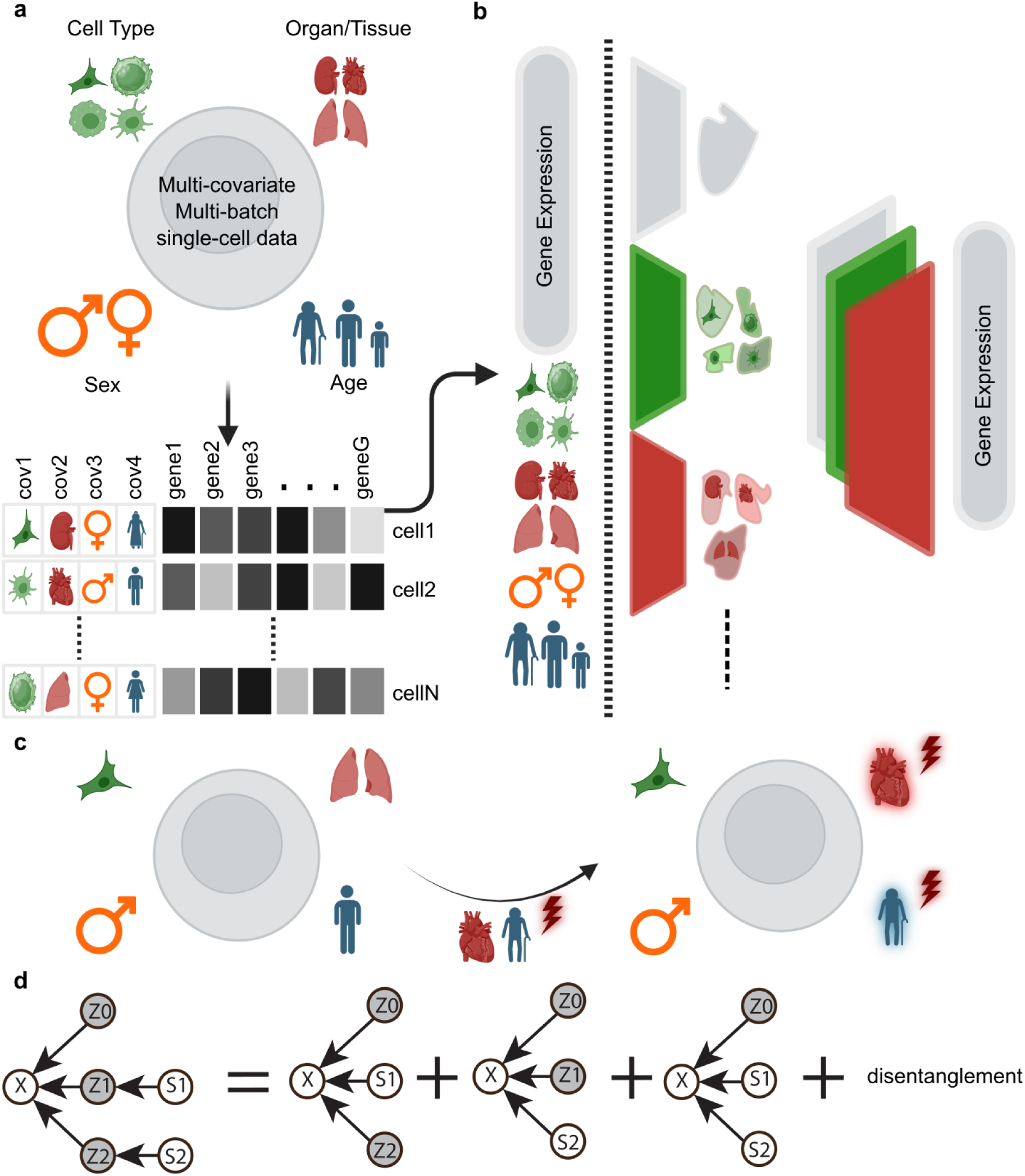
CellDISECT overall architecture to disentangle biological covariates and enable counterfactual predictions. a) CellDISECT provides a multi-facetted interpretation of multi-condition, multi-batch datasets by capturing effects of covariates such as cell type, tissue, sex and age on each cells expression profile. b) CellDISECT is a mixture of expert variational autoencoders (VAEs), one for each covariate plus a VAE for unobserved covariates. Each VAE learns a latent space aligned to its associated covariate but also captures the underlying biology. c) Using a structural causal model, we can generate counterfactual profiles for single-cell data by changing one or more of the values of the covariates. d) A cartoon that explains the mathematical foundation of our method. Our latent spaces are defined by the causal model on the left-hand side, where x is the observed gene expression, S are the observed covariates, and Z are the latent variables. Each expert VAE is derived from this causal model by integrating out a different set of the latent variables and auxiliary neural networks are added to disentangle them.

CellDISECT estimates for cell n the conditional distribution, *p*(*x*_*n*_ | *s*_1*n*_, …, *s*_*Cn*_) in which *x*_*n*_ is the G-dimensional vector of observed RNA counts (G is the total number of genes), and *s*_1*n*_,…, *s*_*Cn*_ are covariate vectors describing information such as batch index of cell, age of donor, tissue etc. In total there are N cells, and C covariates for each cell. This distribution is estimated using a Mixture of C + 1 Expert (MoE) VAEs, and C(C + 1) auxiliary MLPs that decouple these expert VAEs. In addition to interpretability, and flexible batch correction, this decoupling also offers a straightforward approach to counterfactual prediction^15^.

The architecture of CellDISECT is mathematically derived from rigorous manipulations of the stipulated causality graph where we integrate out a different subset of covariates each time (see Supplementary Note 1 for the theory of the model). The generality of our approach and its use of a mixture of disentangled experts also offers three features unique to our approach: 1) supervised latent representations of cells, which capture covariate information and at the same time 2) new biology, thereby providing users with a multifaceted view of the data and facilitating exploration of cell type specificity; 3) flexible fairness in batch correction^21^, which enables the user to define which covariates to use as biological and which as batch at *test time*, as opposed to only at train time as with most methods.

Leveraging both causal inference and disentanglement (see **Figure 1d**), allows CellDISECT to outperform both Biolord and scDisInFact across several tasks in disentanglement and out-of-distribution (OOD) counterfactual predictions, which we benchmark using famous as well as novel metrics. We implemented CellDISECT using the scvi-tools library and made it publicly available to download^22^.

#### CellDISECT captures covariate-specific information in complex single-cell studies

To assess how CellDISECT’s covariate-supervised cell embeddings can capture biological information we applied CellDISECT to the cross-organ dataset^23^ which consists of >200k nuclei of 25 samples from 16 donors. The covariates for each cell in this dataset include the sample ID, organ location, and the donor’s gender and age. The results, presented in **Figure 2**, highlight the novelty of CellDISECT which offers latent spaces that not only capture covariate information but also biological variation. We compare CellDISECT to other baseline models^13,14^ across several metrics, illustrating CellDISECT’s advantages in terms of both latent space structure and quantitative performance.

**Figure 2.**
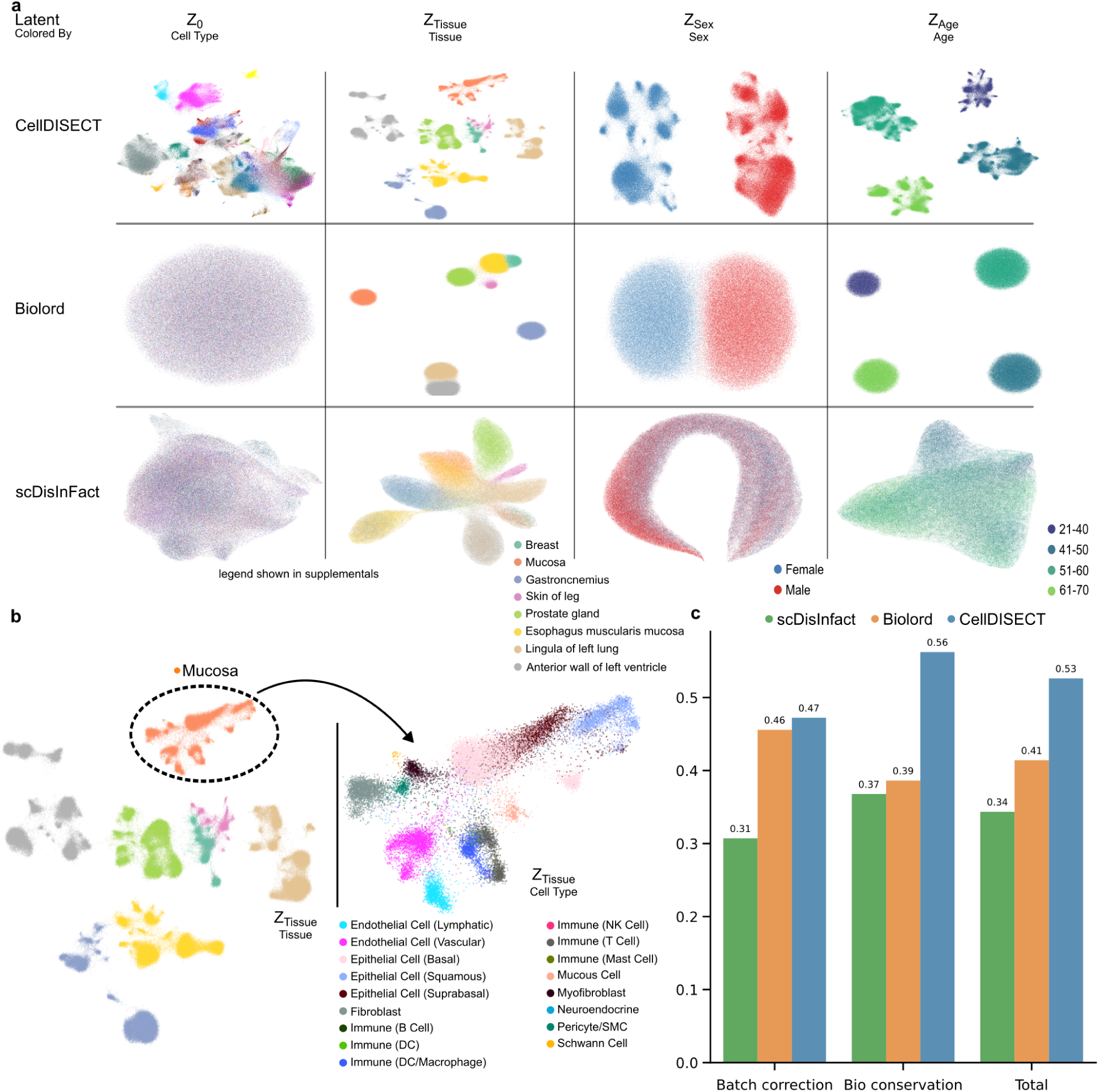
CellDISECT captures covariate-specific information in complex single-cell studies: a) UMAP visualizations of covariate-supervised and unsupervised latent spaces by CellDISECT, Biolord, and scDisInFact on a cross-tissue multi-donor atlas^23^. The supervised cell embeddings in Biolord and scDisInFact only capture the covariate information and are analogous to covariate embeddings. In contrast, CellDISECT provides cell embeddings that can reveal more cell type specificity. b) UMAP visualization of the latent space Z_Tissue_ colored by tissue (left) and magnified portion of mucosa colored by cell type (right). The magnified portion highlights CellDISECT’s ability to distinguish cell types in all its latent spaces even when cell type information is not provided to the model, thereby offering a multifaceted view of the data. c) scIB metrics are used to evaluate the performance of methods in data integration. Here we report for three different methods the average scores of scIB metrics over all given covariates in their respective latent space.

To assess the ability of each model to disentangle covariates, we visualized the latent spaces for CellDISECT, Biolord, and scDisInFact, each coloured by different covariates (**Figure 2a**). The latent space of CellDISECT (top row) clearly clusters cells by each covariate in a structured manner, demonstrating effective separation of biological signals, such as age, sex and tissue. In contrast, supervised cell embeddings generated by Biolord and scDisInFact are akin to covariate embeddings that display more overlap across clusters, suggesting both weaker disentanglement and inability to capture non-covariate biological information within the supervised latent spaces.

In tissue-aligned latent space (see **Figure 2b**), CellDISECT successfully identifies various cell types (e.g., epithelial, immune, endothelial) as distinct clusters, whereas Biolord and scDisInFact do not learn any more biology than tissue specific information. The results suggest that CellDISECT can uncover cell type specificity in an unsupervised manner in all its latent spaces, thereby offering a multifaceted view of the data.

To quantitatively verify the above qualitative observations we evaluate CellDISECT’s integration performance using scIB^24^ (see **Figure 2c**), where we assess both biological conservation and batch correction metrics over all latents, across CellDISECT, Biolord, and scDisInFact. CellDISECT achieves superior scores in both categories, with an aggregate score of 0.53, outperforming Biolord and scDisInFact (aggregate scores of 0.41 and 0.34, respectively). For each covariate given to the model, we ran scIB metrics on Z_0_ and Z_i_ (covariate’s respective latent) with label_key set to the cell type annotations (not an observed covariate), and batch_key set to all other covariates. Showing the ability to discover cell types (Bio conservation), while removing observed covariates’ effects (Batch correction) from all latent spaces (see **Figure 2c** and Supplemental Figures 8-12).

These results demonstrate that CellDISECT effectively balances the removal of batch effects while maintaining the biological relevance of cell types, underscoring its suitability for integrating multi-batch, multi-condition single-cell datasets.

### CellDISECT accurately disentangles covariate information and predicts responses to inflammatory and infectious perturbations

Having verified CellDISECT’s novel ability to offer multi-faceted cell embeddings, we then seek to demonstrate the second use case of CellDISECT: counterfactual predictions with new state-of-the-art performance on biologically relevant perturbations. To benchmark our approach we devised several different scenarios for out-of-distribution counterfactual predictions (see **Figure 3a)**. Importantly we also introduced a benchmark on double counterfactual prediction which to our knowledge marks the first time double counterfactuals are used to benchmark cell perturbations.

**Figure 3.**
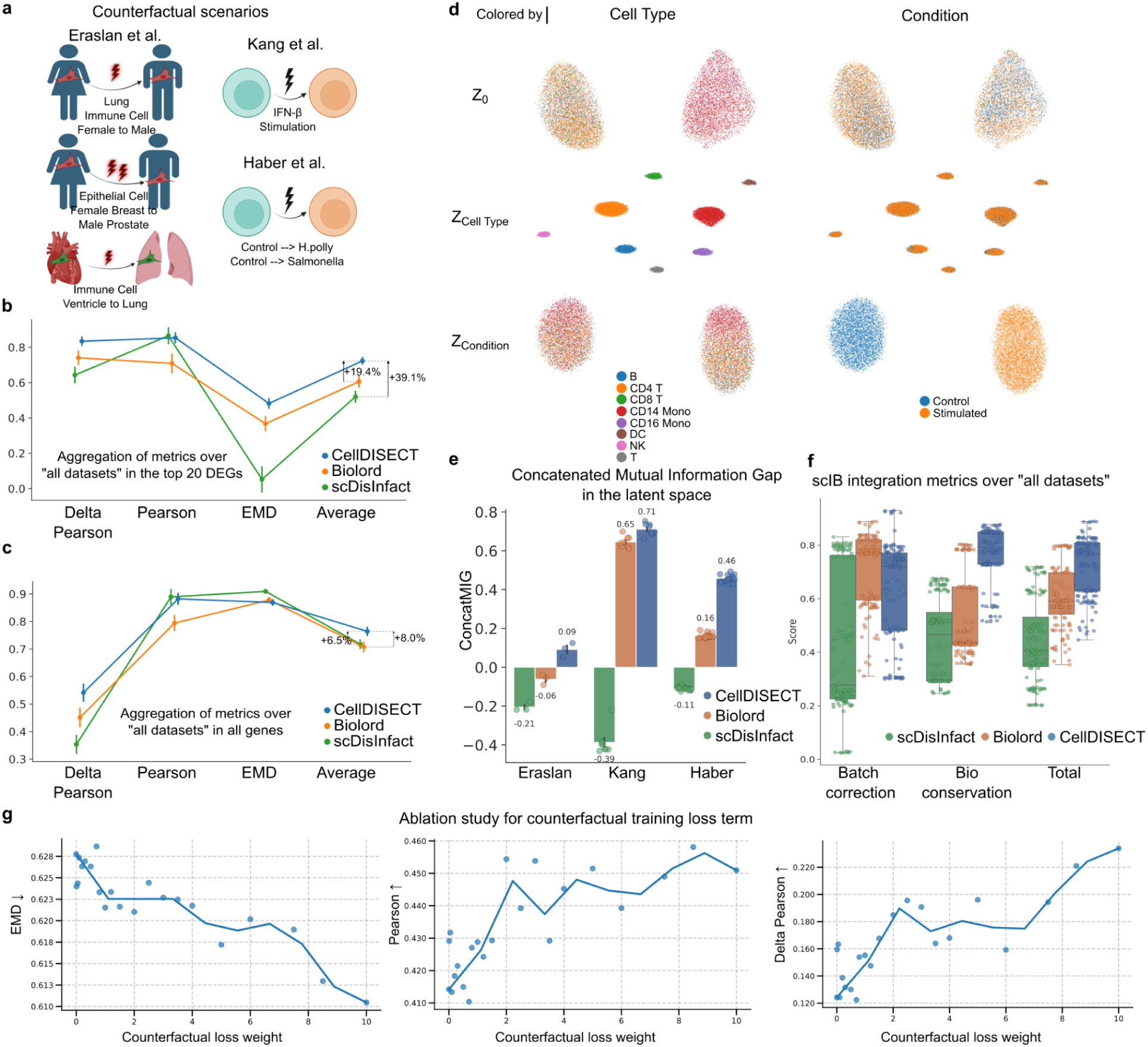
Benchmarks on modelling and disentangling cell type specific response to inflammation, disease and tissue perturbations: a) Illustration of counterfactual prediction scenarios for 3 different datasets used to benchmark our method (for details see Methods section). b) Aggregation of 3 different metrics and their average to evaluate counterfactual prediction capabilities for the top 20 differentially expressed genes. The reported Pearson correlation is between in-silico perturbed and in-vivo perturbed cells across genes. The Delta Pearson reported is the correlation between in-silico perturbed and in-vivo perturbed after the unperturbed gene expression profile has been subtracted from both. The Earth Mover’s Distance (EMD) is the average of distances between in-silico perturbed and in-vivo perturbed cells in gene space. To have all metrics be such that higher value is better prediction, we plot here 1-EMD, instead of EMD (for details see Methods section). c) Aggregation of 3 different metrics and their average to evaluate counterfactual prediction capabilities for all genes. d) UMAP representations of the latent spaces obtained from Kang et al. offer a visualization of CellDISECT’s disentanglement abilities. Each latent space has been aligned to its supervising covariate as evidenced by the disentangled UMAPs along the main diagonal, but is agnostic to the other covariate. We also observe that the unsupervised covariate space uncovered a “lineage” covariate and has separated the cell according to their lymphoid or myeloid lineages. e) Disentanglement comparison of CellDISECT and other models, over 3 different datasets using a novel information theoretic measure, called Concatenated Mutual Information Gap (concatMIG), that captures how much more information a latent space has about its supervising covariate than all the other covariates combined. Higher concatMIG value indicates higher disentanglement. f) Disentanglement comparison of CellDISECT and other models using biologically motivated metrics. We report the integration metrics computed using scIB averaged over all latents of all datasets. g) CellDISECT performance on counterfactual predictions at test time on our counterfactual scenarios for the blood and intestine datasets. The y-axis includes three different metrics: Earth-Movers-Distance (EMD) – where lower is better –, Pearson correlation and Delta Pearson correlation and we report the average of these metrics across datasets, counterfactual scenarios, and gene sets. The x-axis shows the value of the coefficient multiplying the counterfactual term, which is a hyperparameter of our model.

#### Description of datasets

To achieve comprehensive benchmarking for disentanglement and counterfactual prediction we use not only the Cross-organ dataset (introduced above) but also Blood, and Intestine datasets. The Blood dataset^25^ comprises 13,576 cells from control and interferon (IFN)-β stimulated cells, which we treat as a binary covariate, and a categorical covariate indicating the cell type. Finally, the intestine data^26,27^ contains 9,842 individual epithelial cells from the small intestine of mice whose condition is either healthy, infected by Salmonella or by Heligmosomoides polygyrus (H.Poly). For this dataset we used the covariates for “batch”, “condition” and “cell label”.

To benchmark CellDISECT in the Blood dataset, which is a popular dataset used for benchmarking perturbations^3,28^, we used 8 different train/validation/OOD splits, corresponding to the 8 different cell types in the data where in each split the stimulated cells in the corresponding cell type are left out as out-of-distribution. At inference time, and for every split (cell type), we are predicting what the gene expression of these control cell types would have been, had they been stimulated, and comparing it to the held out stimulated cells. We observe that CellDISECT has the highest performance in 5 out of 6 comparisons with Biolord and scDisInFact. These comparisons include three different metrics (Pearson, Delta Pearson and Earth Mover’s Distance (EMD) and two different gene sets (for all genes and for the top 20 differentially expressed genes (DEGs)), see Supplemental Figure 1 and 2.

In the Intestine dataset, we challenged our model even more by performing perturbations from source to target cells, where both the source and the target cells are out-of-distribution (OOD) during training. In particular, for each of its 8 cell types, we constructed 2 scenarios, one with control cells and Salmonella-infected cells being held out, and one with control cells and 10 days H. poly-infected cells being held out (making up to 16 splits). For the counterfactual prediction task in each split, we predict what the gene expression of the infected cells in the corresponding cell type would be, given the control cells. We observe that CellDISECT has the highest performance in 4 out of the 6 comparisons, see Supplemental Figure 3 and 4.

Finally to showcase CellDISECT’s ability to model and perturb many covariates on the same dataset, we devised several perturbation scenarios for the Cross-organ dataset, including a double counterfactual where two covariates are perturbed at the same time. To our knowledge, this is the first time double perturbations are used to benchmark perturbation models for single cell data.

In scenario A, we studied sex perturbation, by holding out (across all organs) all male cells of cell type T, NK, and DC/macrophage; as well as all female cells of these cell types in the lung. During evaluation, we performed counterfactual predictions on these cell types, to predict what the gene expression would be if female cells in the lung were male. In scenario B we devise a double perturbation experiment, where we perturb both the sex of the donor and the tissue environment of the cells while remaining within the epithelial luminal cell type. In particular we hold out all the epithelial luminal cells in the dataset, and then try to predict the male prostate epithelial cell from the female breast epithelial cells. In scenario C, we studied the effect of perturbations on the tissue environment of a cell. We held out all the male cells in the dataset, and trained solely on female cells. Then, we predicted the response of male muscle macrophages upon changing their tissue environment to the lingua of the left lung and then compared to the male alveolar macrophages. In this more complex dataset, CellDISECT exhibited strong performance increases across all 6 comparisons to competing models, see Supplemental Figure 5 and 6.

#### CellDISECT achieves superior counterfactual prediction performance

Benchmarking CellDISECT against Biolord and scDisInFact on the datasets and scenarios, metrics and gene sets we described above, we observe that it exhibits a nearly 20% performance increase in predicting the top20 DEGs (see **Figure 3b**) and 6.5% increase in predicting all genes (see **Figure 3c**), with respect to Biolord, which is the previous state of the art in our benchmarking.

As noted in a recent publication^28^, Pearson correlation becomes a less and less useful metric for comparing models’ performance as their performance improves. Instead, they recommend using Delta Pearson^28^ as a harder and more reliable task when models’ performance approaches pearson correlation of 1. In line with these observations, we report that CellDISECT is roughly tied with Biolord and scDisInfact when benchmarked with the Pearson correlation, but exhibits strong performance increases in Delta Pearson correlation, as well as on average over all metrics. Notably, overall CellDISECT shows strong performance increases in the hard tasks, such as the double counterfactual scenario in Eraslan, where our model predicts the target cells with a Pearson correlation of 0.89, whereas both the competitors and the control cells have negative correlations with the target cells (see Supplemental Figure 7).

#### CellDISECT achieves superior disentanglement of covariates

We can get an intuition of CellDISECT’s approach to disentanglement from the UMAP visualization of its latent spaces on the Kang dataset, see **Figure 3d**. *Z*_*cell type*_ has learned to disentangle cell type information, but is completely uninformative to the condition covariate, conversely, *Z*_*condition*_ disentangles the condition covariate but is uninformative to the cell types. *Z*_0_ has forgotten both the cell type information and the condition information, but instead has learnt another unmeasured covariate, which is the lineage of the cells. Specifically *Z*_0_ clusters all the lymphoid cell types together and separately from all the myeloid cell types, showcasing how CellDISECT can learn new biology in an unsupervised fashion.

We now proceed to quantitatively benchmark the disentanglement properties of CellDISECT against those of Biolord and scDisInFact using information-theoretic (see **Figure 3e**) and biological metrics (see **Figure 3f**).

To benchmark the information-theoretic disentanglement properties of CellDISECT, we introduce a new metric, which we call concatenated mutual information gap (concatMIG), to overcome limitations of MIG ^5^ related to nested covariates. concatMIG calculates the difference (“gap”) of the mutual information between *Z*_*i*_ and covariate i minus the mutual information between all other latents spaces together (*Z*_-*i*_) and covariate i. Across all three datasets, CellDISECT shows the highest level of information-theoretic disentanglement. Notably in the Cross-organ dataset, not only CellDISECT exhibits superior disentanglement, but the competing methods report a negative concatMIG which implies that all the non-aligned latent spaces when taken together contain more information about a covariate than the covariate space which is aligned to it.

Moreover, to benchmark the biological disentanglement properties of CellDISECT we employ the SCIB metrics, a widely used framework for evaluating methods on tasks of batch effect correction and biological information preservation. For the Blood and the Intestine dataset, we applied scIB on each of the latents *Z*_*i*_ s.t. i > 0 with the label key (the covariate we expect the latent space to be preserving; bio conservation) set to the covariate aligned to it, while setting the rest of the covariates (once at a time) to the batch key (the covariate we expect the latent space to be ignorant of). As for the Cross-organ dataset, since the cell type is not provided as a seen covariate while training, we are calculating the scIB metrics over each of the latents (including *Z*_0_) with the label key set to the cell type annotations from the dataset, and the batch key set to all the seen covariates, that are non-correspondent to the latent each time. The final values reported in **Figure 3f** show the aggregated mean values of scIB metrics on all 3 datasets.

Finally, to test the effect that causal analysis on synthetic counterfactuals *during* training has on CellDISECT’s performance on counterfactual predictions at *test* time we perform an ablation study of the counterfactual term in the loss function. In our model we introduce hyperparameters as coefficients of all the terms in the loss function. In our ablation experiment, we keep the coefficients of all the terms in the ELBO as derived in proposition 1 (see Methods) and vary the coefficient associated with the counterfactual term. We observe that ablating this counterfactual loss term leads to reduced model performance in counterfactual predictions at test time (see **Figure 3g**). In more detail, CellDISECT’s performance deteriorates without the counterfactual term in 9 out of 12 metrics, stays roughly constant in 1 and improves only in 2 (see Supplemental Figures 15, 16).

### CellDISECT identifies non-classical megakaryocyte sub-population

We now demonstrate how CellDISECT’s latent spaces, which capture covariate information as well as biological variation, can be used to explore cell type specificity. In particular, CellDISECT identifies a new subpopulation of megakaryocytes which exhibits immune behavior.

#### Description of data

A recent study^18^ mapped the developing human immune system across nine prenatal organs using single-cell RNA sequencing and spatial transcriptomics, creating an atlas of over 900,000 immune cells, see **Figure 4a**. They discovered system-wide blood and immune cell development beyond traditional haematopoietic sites, including the gut and skin. The study highlighted the maturation of monocytes and T cells before migration to peripheral tissues, characterised innate-like B1 cells and unconventional T cells, and identified lineage-committed progenitors in unexpected locations.

**Figure 4.**
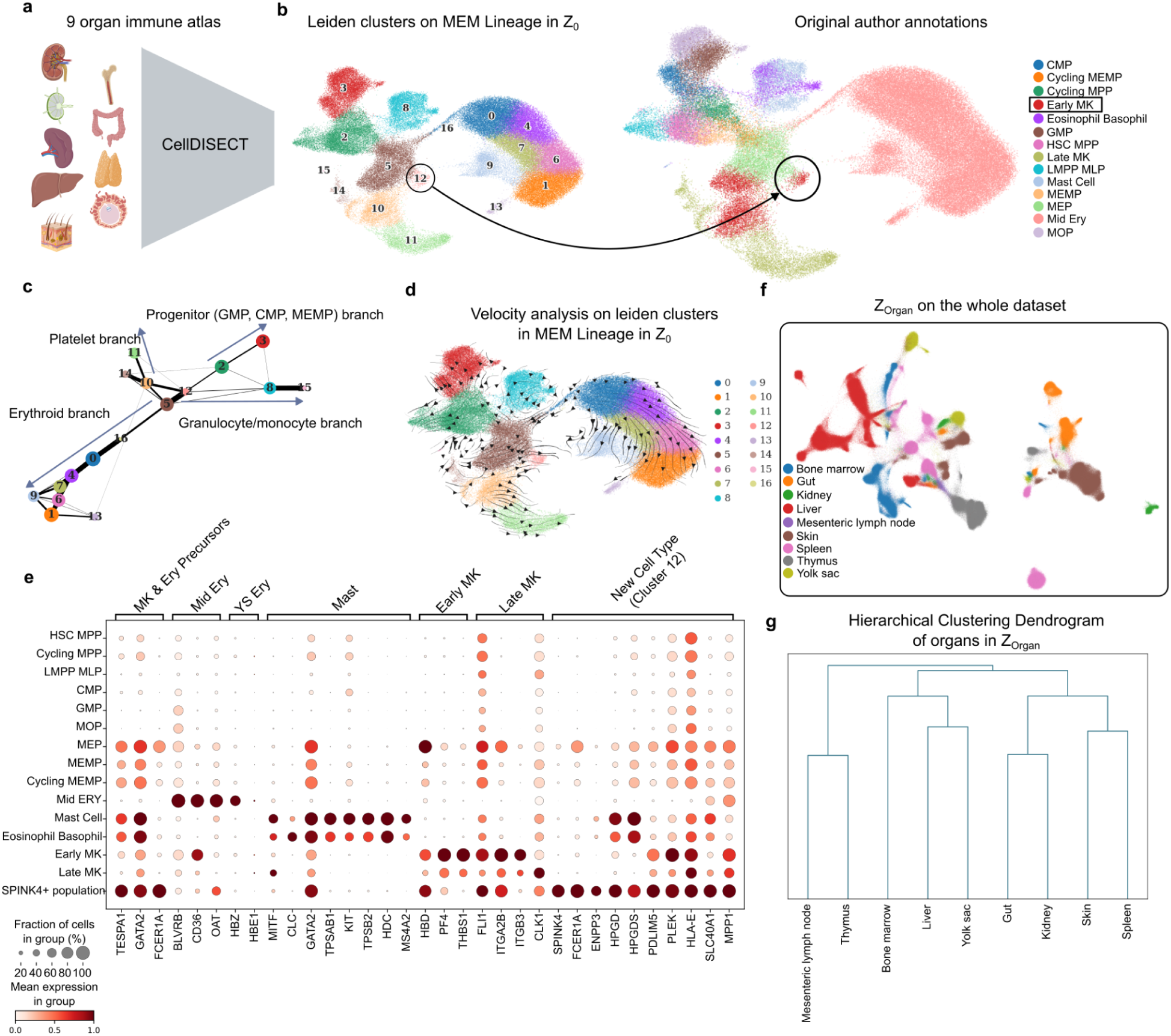
CellDISECT identifies a non-classical megakaryocyte subpopulation. a) Schematic application of CellDISECT on the 9 organ developing immune system atlas^18^ and UMAP visualization of the megakaryocyte-erythroid-mast (MEM) lineage in CellDISECT’s unsupervised latent space, *Z*_0_. The UMAP is colored by Leiden clustering in CellDISECT’s *Z*_0_ latent space identifies a (non-classical) hematopoietic subpopulation annotated as an early megakaryocyte in the original publication^18^ but distinct from other cells annotated as early megakaryocytes. c) Connectivity analysis of the latent manifold using PAGA reveals the new population to be located close to MEP and early MK populations. d) Trajectory analysis using sc-velo shows that the non-classical MK subpopulation is moving away from MEM but not in the direction of platelet or granulocyte differentiation. E) Dotplot of clusters as originally annotated^18^ against relevant gene markers. The non-classical (SPINK4+) subpopulation expresses known transcription factors active in MK cells, such as FLI-1, but also TFs and gene markers active in mast cells, such as MITF, HPGD, HPGDS, and does not express THBS1 and has reduced expression of PF4 which are known markers that drive differentiation into platelets. It also expresses unique gene markers such as SPINK4, whose expression has not been previously reported in hematopoiesis. f) UMAP visualization of the MEM lineage in the latent space *Z*_*organ*_ which is supervised by the organ covariate. *Z*_*organ*_ learned a biological manifold stratified by organ. g) The stratification of *Z*_*organ*_ by organ covariate is biologically meaningful with more similar organs located closer together.

#### Megakaryocyte biology

Megakaryocytes (MK) are large bone marrow cells responsible for producing platelets, which are critical for blood clotting. They arise from megakaryocyte-erythroid progenitors (MEPs) and undergo a unique maturation process called endomitosis^29^, where they replicate DNA without dividing, leading to their large size which is where their name derives from. Mature megakaryocytes then release platelets into the bloodstream by shedding cytoplasmic fragments^29^. Previous studies^19,30,31^ have shown the possibility for immune function of such cells. One study^19^ showed MK cells in the yolk sac (YS) using single-cell transcriptomics and identified two MK subpopulations with likely niche support and immune functions respectively, suggesting potential immune-like functions for MK cells in primitive haematopoiesis. Additionally, studies in mice^31^ and adult bone marrow^30^ cells also show the potential for immune-like MK cells in adult life.

#### SPINK4 MK subpopulation in developing immune system

We processed the immune atlas^18^ using CellDISECT with default settings. Using Leiden clustering (with default scanpy settings) in the *Z*_0_ latent space in the CellDISECT embeddings (see **Figure 4a**) we observe that the cells annotated as early MK cells in the original publication^18^ split into two distinct clusters (see **Figure 4b**). The smaller of the two clusters contains mainly contributions from spleen, liver, and bone marrow from different donors (see Supplementary Figure 14). Connectivity and trajectory analysis (see **Figure 4c,d**) shows SPINK4+ early MK cells differentiation away from MEP into the MK lineage but not following into late MK cells. This subpopulation expresses MK gene markers (see **Figure 4e**) such as FLI1^32,33^, which is a known key MK lineage TF, and PLEK^34^. At the same time, however, these MK cells do have low expression of PF4 and lacking expression of THBS1 indicating that they are not moving towards platelet formation. They also express MITF, a known mast cell TF, and genes related to mast cell/basophil functions including FCER1A and ENPP3^35,36^, suggesting a potential immune function. They also express HPGD and HPGDS^37^, enzymes involved in prostaglandin synthesis released by activated mast cells that mediate inflammatory functions. However, the differentiation trajectory makes it clear that they are somewhere between MEP and MK in the lineage, and therefore not along the mast cell differentiation trajectory. This suggests that these MK cells may share immune-like roles similar to other immune cell types, showing post primitive haematopoiesis MK cells can also exhibit immune roles during definitive haematopoiesis. The newly identified subpopulation also expresses unique gene markers such as SPINK4 whose function is not yet clear. SPINK4 is primarily linked to goblet cell function ^38–40^ but expression in the hematopoietic lineage has not been previously reported. It is worth noting that Leiden clustering with default settings (of the same cells) in the scVI latent does not assign this new subpopulation to a separate cluster (see supplemental Figure 13) which highlight CellDISECT’s ability to resolve distinct clusters that are masked by scVI embeddings.

Finally, the *Z*_*organ*_ latent space provides a latent representation of the immune atlas stratified by organ (see **Figure 4f**). The stratification of the organs in this space captures the underlying biology between organs, with more similar organs located more closely (see **Figure 4g**). Bone marrow, yolk sac, and fetal liver are all major hematopoietic organs, whereas lymph nodes and thymus are major lymphoid organs rich in lymphocytes such as T cells.

In conclusion, we identify a subpopulation of early MK cells during fetal development which exhibits immune-like roles during definitive haematopoiesis, distinct from MK cells with immune-like roles previously reported in the YS.

### CellDISECT highlights erythropoiesis difference between liver and bone marrow by leveraging multiple latents

We now demonstrate how CellDISECT’s multiple latents not only offer a multifaceted view of the data but at the same time enable flexible fairness in single-cell integration. CellDISECT allows selecting which covariates to designate as biological and which as batch covariates at test time, as opposed to only at train time like other methods. We then leverage this ability to highlight differences in erythropoiesis between the liver and bone marrow.

#### Erythropoiesis biology

Erythropoiesis is a highly regulated, key function which is transferred between organs throughout embryonic and fetal development^41^. Two of the key organs this occurs in is the liver, the primary haematopoietic organ during development, and the bone marrow which is then the main organ responsible for erythropoiesis throughout adulthood.

#### Differences in erythropoiesis between liver and bone marrow

To study erythropoiesis in the ^18^ atlas, we leverage the flexible fairness inherent in our method by concatenating the latent representations from *Z*_0_ and *Z*_*organ*_ to construct the *Z*_0,*organ*_. This latent space is aligned with both the organ covariate and unknown covariates, and has the effect of the platform and donor covariates regressed out (see **Figure 5a**). Notably to construct this latent space, we did not need to retrain our model, but manage to construct it at test time by leveraging the flexible fairness property of our method. Subsetting the dataset to the erythropoiesis branch, as annotated in the original publication^18^ (see **Figure 5b**), we observed that erythroids would split into two main routes caused by the biological covariate organ; depending on if they are contributed by the liver or the bone marrow, and not due to technical effects. Other organs would also split however, with the low number of erythroid contributions from these organs however, these differences are less prominent due to the lack of cell numbers.

**Figure 5:**
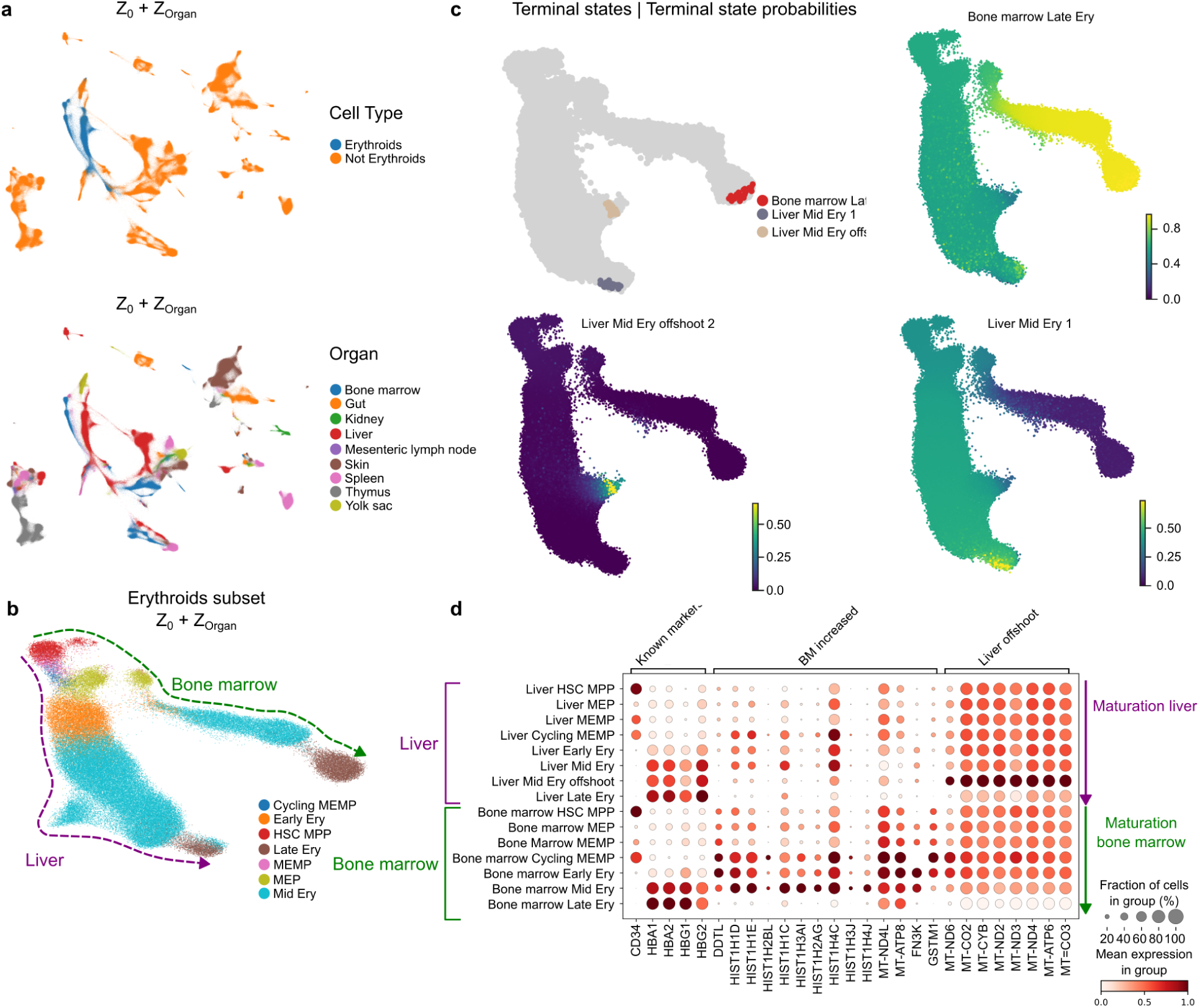
CellDISECT identifies erythroid maturation difference between liver and bone marrow. a) UMAP visualisation of Erythroids (top) and organ (bottom) distribution in *Z*_0,*organ*_ derived embedding. b) UMAP visualisation of Erythroid lineage plotting original author cell types. Arrows indicating direction of maturation for liver (purple) and bone marrow (green). c) Terminal states for erythroid maturation determined from GPCCA model using cytotrace kernel from Cellrank. Left showing 3 terminal cell states. Right showing fate probabilities of all cells for each terminal cell state identified. d) Dot plot showing the mean expression (colour) and proportion of cells expressing proteins (dot size) of known erythroid maturation markers, markers separating Liver and Bone marrow lineages being increased in Bone marrow and Mitochondrial markers separating liver specific erythroid cluster offshoot

To investigate these 2 splits of erythroid maturation seen between the liver and bone marrow, trajectory analysis was performed using Cellrank on *Z*_0,*organ*_ (see **Figure 5c**). This analysis shows that the split between organs first becomes apparent at the MEP stage with liver erythroids further splitting at mid-erythroid stage primarily linked to increased mitochondrial activity. Gene trends of known erythroid maturation markers (HBA1, HBA2, HBG1, HBG2, HBB) shows both routes increasing across as well as highlighting bone marrow erythroids having increased expression of histone genes compared to liver, occurring in the early to mid erythroids (see **Figure 5d**). As erythroid maturation is modulated through histone regulation by processes such as acetylation, methylation and phosphorylation^41–43^, this gene expression pattern suggests a higher proportion of histones may be produced in bone marrow erythroids. An increase in histone levels and accessibility could enable greater control over erythroid maturation providing more sites for control which could be one contributing reason why the bone marrow takes over the role of erythropoiesis from the liver and into adulthood.

In summary, CellDISECT’s flexible fairness enables users to identify and explore gene expression patterning linked to a given category of interest. Without a need to retrain, CellDISECT offers many supervised latent spaces, some of which provide negative controls where the category of interest is regressed out by treating it as a batch covariate. This multifaceted view afforded by CellDISECT enhances data exploration and interpretability relative to other methods.

## Discussion

CellDISECT represents a significant advancement in single-cell analysis by integrating disentangled representation learning with causal analysis on synthetic training data, and addresses the complexity of multi-batch, multi-covariate data. Its novel architecture and use of synthetic data during training enables covariate-specific latent spaces while retaining the ability to capture biologically meaningful variations. This dual capability not only enhances interpretability but also allows for robust counterfactual predictions, offering a new dimension in single-cell research.

Compared to existing methods such as Biolord and scDisInFact, CellDISECT demonstrates superior disentanglement and predictive accuracy, particularly in out-of-distribution scenarios.

CellDISECT is able to model both observed and unobserved covariates while maintaining biological relevance, enabling discovery of cell types at enhanced resolution. For instance, its application to the immune atlas uncovered a previously unidentified megakaryocyte subpopulation with immune-like characteristics.

Although it has been shown that MK cells can have an immune role in the YS and in adult life, its entire immune repertoire is yet underexplored. We highlight the identification of immune-like MK cells during early human development in the spleen, liver and bone marrow across multiple donors at single cell resolution. Unlike classical platelet-producing MKs, these cells exhibit high expression of immune-related genes, such as FCER1A, ENPP3 and HPGDS, suggesting potential roles in immune modulation, as well as the unique marker SPINK4. This discovery underscores the potential of CellDISECT to unveil novel cell populations.

CellDISECT also introduces flexibility in defining biological and batch covariates, addressing a limitation of traditional methods that require such decisions during training. This adaptability makes it a powerful tool for integrating heterogeneous datasets, as demonstrated in its ability to resolve complex tissue-specific and donor-specific patterns in cross-organ single-cell data.

Despite its strengths, CellDISECT’s reliance on pre-defined covariates may limit its applicability to datasets with unknown covariate structures. Future developments could explore adding more unsupervised latent spaces for inferring latent variables directly from the data.

In conclusion, CellDISECT establishes a new benchmark for single-cell analysis, combining interpretability, biological discovery, and predictive power. Its ability to disentangle and leverage covariates paves the way for deeper insights into cellular heterogeneity and its drivers.

## Data Availability

The real data can be downloaded from the links. We downloaded the annotated dataset^23^ in h5ad format from the CZI CELLxGENE dataset hub^44^. The blood dataset^25^ was downloaded and obtained from CPA^3^ tutorials. The anndata object for the mouse intestine dataset^26^ was obtained using the Pertpy library^45^. The dataset for the developing immune system^18^ was provided by the original authors but is also publicly available^46^.

## Code Availability

Our source code for the method is available at https://github.com/Lotfollahi-lab/CellDISECT and the code for reproducing the results of our experiments is available at https://github.com/stathismegas/CellDISECT_reproducibility.

## Acknowledgments

We would like to thank Dr. Arash Mehrjou for helpful feedback and discussions. This work was made possible in part by the Wellcome Trust (220540/Z/20/A) and the Accelerate program for Scientific Discovery. M.H. is funded by Wellcome (WT107931/Z/15/Z, WT223092/Z/21/Z, WT206194, and WT220540/Z/20/A), the Lister Institute for Preventive Medicine, and NIHR and Newcastle Biomedical Research Centre. The funders had no role in study design, data collection and analysis, decision to publish, or preparation of the manuscript. Figure cartoons were created with BioRender.com.

## Author Contributions

S.M. conceived the theory of the model and the perturbation modeling, implemented it in code and as a GitHub package, and wrote and edited the manuscript. A.A. performed the benchmarking on simulations and real data, applied the model on the immune atlas, improved the code implementation and documentation, and helped edit the manuscript. A.R. led the biological interpretation of the results on the immune atlas and helped edit the manuscript. O.D. performed literature research for gene markers and helped edit the manuscript. K.S. wrote parts of the code in the early stages of the project. H.A. implemented the continuous embedding of the categorical covariates. K.P. helped create conda environments for the project. M.H. provided key biological interpretation, and supervised the project. S.A.T. provided key biological interpretation, and supervised the project. M.L. conceived the idea of disentangled representations in single-cell data and of a model that provides multiple latent spaces for each covariate, and supervised the project.

## Methods

### Datasets

#### Collection and pre-processing

The datasets, their count matrices, cell types, and associated metadata in this study were downloaded from the original study or from CellxGene data portals where the authors have submitted them. All the preprocessing steps such as gene filtering, log normalisation and highly variable genes (HVG) selection were performed using scanpy^47^) unless stated otherwise.

*Eraslan dataset*^23^ was downloaded from CZ CELLxGENE^48^ and count matrix comprised of 209,126 cells and 32,922 genes. Genes that have at least 3 counts and log-normalised data was used to select 1212 HVGs comprising 1200 HVGs (highly variable genes), and “Toll-like receptor” genes were used in the model.

*Kang dataset*^25^ was downloaded from CPA^3^ tutorials, preprocessed, filtered, and 5000 HVGs were used in the model. The dataset consists of 13576 PBMCs, from eight lupus patients. These blood cells are either untreated or treated using IFN-β stimulation. (7217 stimulated and 6359 control cells)

*Haber dataset*^26^ was downloaded from the Pertpy library^45^ using the command “pertpy.data.haber_2017_regions()”. We filtered the dataset to genes that are expressed in at least 20 cells, log-normalized the counts, and selected 7000 HVGs for training the model. This dataset consists of intestinal epithelial cells of mice, after being subjected to Salmonella (1770 cells) or Heligmosomoides polygyrus (H. poly) infections (2121 after 3 days and 2711 after 10 days), and 3240 control cells.

*Suo dataset*^18^ was provided by original authors and then subsetted to retain only good quality cells. Good quality cells were determined by selecting only cells which were used to generate part of the Human_Whole_Embryo/v1 model from CellTypist^46^. This model compared cells against multiple other public fetal datasets and is publicly available^46^. Original author annotations were then used throughout analysis. The data was log-normalized the counts using Scanpy’s “normalize_total” and “log1p” methods. Finally, we selected the top 8192 HVGs ranked by gene dispersion norm following the same logic as the original publication. As input covariates for CellDISECT to model, we provided donor and chemistry (3GEX/5GEX) matching the original publication, as well as organ.

#### Splits

##### Haber dataset

We constructed 16 different train/validation/ood splits, and the results are reported as the average of these splits. For each cell type (there are 8 cell types in the dataset), we constructed 2 splits, one with control cells and Salmonella-infected cells being held out, and one with control cells and 10 days H. poly-infected cells being held out (making up to 16 splits). For the counterfactual prediction task in each split, we predict what would the gene expression of the infected (infection corresponding to the split) cells in the corresponding cell type be, given the control cells. Both source and target cells are unseen and out-of-distribution in this case.

##### Kang dataset

To achieve comprehensive benchmarking, we used 8 different train/validation/OOD splits, corresponding to the 8 different cell types in the data where each in each split, all the stimulated cells in the corresponding cell type, are left out as out-of-distribution. Accordingly, in each split (cell type), we predict what the gene expression of the stimulated cells would be, given the control cells in that cell type. These splits were available in the downloaded object from CPA^3^ tutorials.

##### Eraslan dataset

To do counterfactual predictions and benchmark our method against other methods in different scenarios, we created 3 different splits in the dataset, each with a different Out-Of-Distribution (OOD) set of cells (Fig. 3a).

Scenario A: Immune T, NK, and DC/macrophage cells were held out from all male cells across organs, as well as within the lung for male cells to acquire an out-of-distribution set. During the evaluation, we performed counterfactual predictions on “Immune (DC/macrophage)” cells, to predict what the gene expression would be if female cells in the lung were male. Scenario B: This scenario’s OOD cells consist of all the “Epithelial cell (luminal)” cells. Then we performed a double counterfactual prediction, evaluating the methods for predicting what would the gene expression for Epithelial cells in the female breast tissue be, if they were actually male prostate gland cells. Scenario C: In this case, we held out all the male cells in the dataset, and trained solely on female cells. Then, we performed counterfactual prediction, on “Male” Immune DC/macrophage and alveolar macrophage cells in the anterior wall of the left ventricle, to predict what their gene expression would be if they were the same cells in the lingula of the left lung.

### Model

#### Theoretical Foundations

We provide here a brief overview of the theory and architecture of CellDISECT, for more details see Supplementary Note 1. CellDISECT estimates for cell n the conditional distribution, *p*(*x_n_* |*s_1n_*, *s_2n_*, …, *s_Cn_*) in which *x*_n_ is the G-dimensional vector of observed RNA counts (G is the total number of genes), and *s_1n_*, …, *s_Cn_* are covariate vectors describing information such as batch index of cell, age of donor, tissue, etc. In total there are N cells and C covariates for each cell. This distribution is estimated using a Mixture of C +1 Expert (MoE) VAEs, and C(C +1) MLPs that (de)couple these expert VAEs. In addition to interpretability, and flexible batch correction, this decoupling also offers a straightforward approach to counterfactual prediction as explained in Foster et al. (2022), and we leverage this by training our algorithm to maximize the sum of the likelihood of the observed data and the likelihood of the model’s counterfactual predictions. Throughout the rest of the paper, when we refer to a specific cell or generally the random variable, we will drop the superscript *n*.

CellDISECT provides users with a multifaceted view of the data with many different latent spaces and VAEs, each aligned with a different covariate. We get each of these expert VAEs by mathematically marginalizing out all covariates to which we don’t want to align, as shown in the cartoon of Figure 1d.

With these specifications the architecture of the encoders and decoders of the model follows mathematically from our structural causal model (SCM), see Supplementary Figure 17. This derivation is the content of Proposition 1 which also derives the appropriate ELBO function of our model.

##### Proposition 1.

The factorization of the SCM in figure 1 implies the following evidence lower bound,

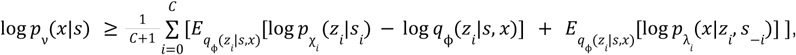

where ν = {χ*_i_*, λ*_i_* : ∀*i*}. As Proposition 1 shows, CellDISECT has C+1 encoders, 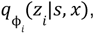 which take as input all the covariates and the gene expression; C+1 decoders, 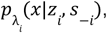 which take as input a latent variable, and all the covariates except for the one corresponding to this latent variable; and C+1 prior encoders, 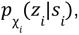 to allow more flexibility in learning the prior distributions for the latent variables, which has also been shown to promote identifiability of the latent space variables^49^.

*Sketch of proof*: We marginalise out all covariates but one and derive an ELBO for the reduced SCM, then average all these ELBOs. For details see Supplementary note 1.

Our approach to implementing counterfactual predictions in Pearl’s three-step approach is inspired by Foster et al^15^, which streamlines it by imposing an orthogonality constraint between latent variables and observed covariates. However, their constraint needs to be generalized to meet the demands of our multi-covariate model. Our proposed generalization is the constraint ∀*i* ≠ *j* : *z_i_* ⊥ *s_j_*, which demands that different representations are aligned to only one covariate at a time. We then follow the three steps in Pearl’s method of counterfactual prediction^6^:

1. abduction: we infer the value of the root nodes {*z*_0_ } based on the observations, *p*(*z*_0_|*s*, *x*),
2. action: swapping *s_i_* for *s_i_* ’, *do*(*s* = *s_i_* ’),
3. prediction, 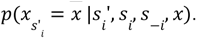

The fundamental problem in causal inference is that we have access only to one outcome (termed factual). We never observe the counterfactual outcome, which is defined by the do operations on the causality graph and therefore might not even be identifiable from the observations we actually performed. As we show in proposition 2, in our model, the counterfactual outcome of intervention on any covariate is identifiable by data that has been observed via functions learned by our model during training.

##### Proposition 2.

The counterfactual (never observed) value of the response variable x due to an intervention *do*(*s*_*i*_ = *s*_*i*_’) on any covariate i can be identified based on the rules of counterfactual reasoning by observed data and is equal to,

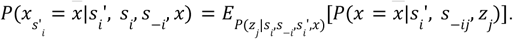

We notice that the RHS contains only ordinary probabilistic expressions (involving no counterfactuals). Moreover, the distributions 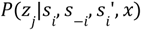 and 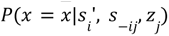 are the j-th encoder and decoder (as derived in proposition 1) respectively, and are learned by CellDISECT during training.

The identifiability of counterfactual predictions allows us to use the idea in Wu et al. (2002)^16^ of semi-autoencoding an algorithm’s counterfactual predictions and jointly maximizing the likelihood of both real data and counterfactuals, log p(x|s) + log p(x′|x, s, s′), where *s*’ = *s_−*i*_* ∪ *s*’*_*i*_*. To implement the counterfactual semi-autoencoding we generalize Wu et al. (2022) to the case of multiple conditions.

Specifically, before training, we generate counterfactual pairs of cells. Every cell c is paired with all other cells c′ of the same cell type that have at most one covariate different. Let *s* be the covariate that differs in the pair (c, c′). Then, during training, we predict counterfactual gene expressions values associated with the new value of *s* (one *s* at a time) and penalize them in the loss function using negative log-likelihood loss under the learned distributions.

#### Model Architecture

##### ENCODERS

CellDISECT employs n + 1 encoders, parameterised by *ϕ*_0_, *ϕ*_1_, …, *ϕ_n_*, to independently infer n + 1 latent vectors *z*_0_, *z*_1_, …, *z_n_*. Each encoder receives the same input: the gene expression vector X and the complete covariate vector s. Additionally, we use n encoders, parameterised by ψ_0_, ψ_1_, …, ψ*_n_*, to infer n prior distributions for the latent variables. The input to the encoder ψ consists solely of *s_k_*. For the latent variable *z_k_*, we apply amortised inference 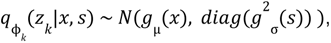 where *N*(µ, σ) is a Normal distribution of mean μ and variance σ. In line with Proposition 1, we infer a prior distribution for *z_k_*as: 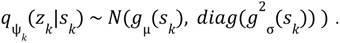

##### DECODERS

We implement n + 1 decoders, parameterised by θ_0_, θ_1_, …, θ*_n_*, to generate the gene expression vector x within each decoder independently. The input for decoder 0 is (*z*_0_; s), while for the k-th decoder (where *k* ≠ 0), the input is (*z_k_*; *s*_-_*_k_*), where *s_k_* refers to all attributes except *s*_-_*_k_*. The rationale behind this setup is to ensure that *z_k_*encapsulates all the information in *s_k_* but none from *s*_-_*_k_*. This architecture enables CellDISECT to learn disentangled representations. Moreover, *z*_0_ is designed to exclude any information about s, instead learning cell-specific background variation due to unobserved covariates. Using decoder θ*_k_*, we predict the expression of gene g using a Zero Inflated Negative Binomial (ZINB) distribution, or alternatively a Negative Binomial (NB) distribution, which has been shown to appropriately model gene expression data Lopez et al. (2018). The gene expression for gene g in cell j is given by: *x_g_* ∼ *NB*(*l f*^*g*^(*z*), *d_g_*), where *l* is the observed random variable representing the number of RNA molecules captured for that gene in the cell. f is a function that maps the latent space to the simplex of the gene expression space, and dg is the dispersion parameter of the negative binomial distribution. By default, we use the ZINB distribution in the generative process.

##### ABLATION STUDY

To benchmark the effect of the counterfactual term on CellDISECT’s performance in counterfactual prediction, we used 22 different values in the range [0,10] for the weight for the counterfactual term and trained CellDISECT models on the Blood and Intestine datasets. We then performed predictions for the different scenarios and splits (see above “Splits”) and evaluated the quality of the predictions using our three metrics for counterfactual predictions: EMD, Pearson correlation, and Delta Pearson. Throughout these experiments we kept the values of most of the rest of the weight hyperparameters equal to 1, which is the value they get assigned in the ELBO. The exception was the prior loss term, whose contribution to the total loss was found to be disproportionately large under this uniform scaling, and was therefore assigned a smaller weight of 0.0003. This adjustment was necessary because, with the default prior weight of 0.003, the prior loss term began to dominate the overall loss function – an effect not observed when using the full set of default hyperparameters – and caused premature early stopping – after 5 epochs – before convergence. Importantly, this rescaling does not reflect a change to the model’s default hyperparameter configuration, but rather ensures stability in the context of this specific ablation study, where other weights were set to 1.

#### Hyperparameter tuning

##### CellDISECT Hyperparameters

To demonstrate the robustness of our method in various tasks, use cases, and datasets, we used the same set of hyperparameters for our experiments across all datasets. We are using encoders/decoders with 2 layers each, 128 hidden neurons, latent dimension 32 for Z_i_ for all i, a dropout rate of 0.1, a learning rate of 0.003, weight decay of 0.00005, and the Adam optimizer.

Biolord hyperparameters: We used the proposed hyperparameters in Biolord’s tutorials and documentations, with two changes after discussions with the method’s authors: first, decreasing the early stopping patience due to the observation of extreme overfitting on the training data in some cases, and second using the “nb” distribution instead of “normal” to acquire counts for better comparison with scDisInFact and CellDISECT.

scDisInfact hyperparameters: We used the proposed learning hyperparameters in scDisInfact’s tutorials and documentation, while increasing the size of the latent dimensions from 8 and 2, to 40 and 10 for the shared and unshared latents respectively, as well as increasing the maximum training epochs to give the model more time and flexibility to learn. These changes were decided and made to keep the comparison fair, due to the observation of scDisInFact achieving better results in the benchmarking using the increased dimensions and epochs.

##### Choice of DEGs in benchmarking

For each of the counterfactual prediction scenarios, we performed a curated computation process to determine the top differentially expressed genes that mark the difference between control and target cells. We used Scanpy’s rank_genes_groups function with the Wilcoxon method to find these genes, within each cell type, between the control and target cells. We ranked these genes by the absolute value of their log fold-change values and took the top 20 DEGs from this ranking to perform evaluations specified for each task and scenario, on all models.

#### Counterfactual training when cell type is not provided

In scenarios where cell type annotations are unavailable, unreliable, or undesired (such as our analysis on the Cross-Organ dataset) CellDISECT is designed to learn and contain the cell type effects as an unobserved covariate within the latent spaces (Fig. 2A, B). This flexibility facilitates exploratory analyses, but introduces a challenge to generating the counterfactual pairs for the evaluation of the counterfactual term. Since there is no cell type annotation given to CellDISECT, counterfactual training which involves pairing cells at random to generate predictions, may pair cells from distinct cell types, which could lead to model collapse. To address this challenge, we implement a preprocessing and augmentation strategy for scenarios where the cell type information is not provided as a seen covariate. Prior to training, Leiden clustering is applied to the PCA-transformed gene expression profiles, partitioning cells into clusters that serve as proxies for cell type groupings. During counterfactual training, cell pairs are sampled exclusively from within the same cluster, ensuring that the counterfactual predictions involve biologically similar cells. This approach prevents the model from encountering mismatched cell types, thereby preserving its capacity to capture cell type-specific information while training.

### Model assessment

#### Metrics

##### Disentanglement

We assess the disentanglement capabilities of the methods using two key metrics. For the first metric, we apply the widely used scIB metrics to evaluate how effectively each latent space preserves its relevant covariate information (biological conservation) while minimizing information related to noise covariates (batch correction).

For the second disentanglement metric, we introduce concatMIG, a variation of the Mutual Information Gap (MIG) metric originally developed by Chen et al. (2018). concatMIG assesses the disentanglement of a covariate from all latent dimensions (spaces) except its corresponding one. To compute this, we compute the difference of the mutual information between each latent and its covariate, minus the mutual information between the concatenated remaining latents and that covariate. concatMIG reports the average of those differences across covariate,

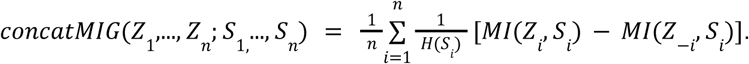

By design, concatMIG can yield negative values; for example, when the non-age latent spaces together contain more information about age than the age-specific latent space does.

##### Counterfactual Prediction

To evaluate and report the counterfactual gene expression prediction abilities of each model, we utilized three different metrics once on all the genes the models were trained on, and once only on the top 20 DEGs (chosen as explained above).

1. Pearson correlation,
2. Delta Pearson Correlation (Pearson correlation between the difference of true minus control expressions and the difference of predicted minus control expressions),
3. Earth Mover’s Distance.

## Supplementals

**Supplementary figure 1.**
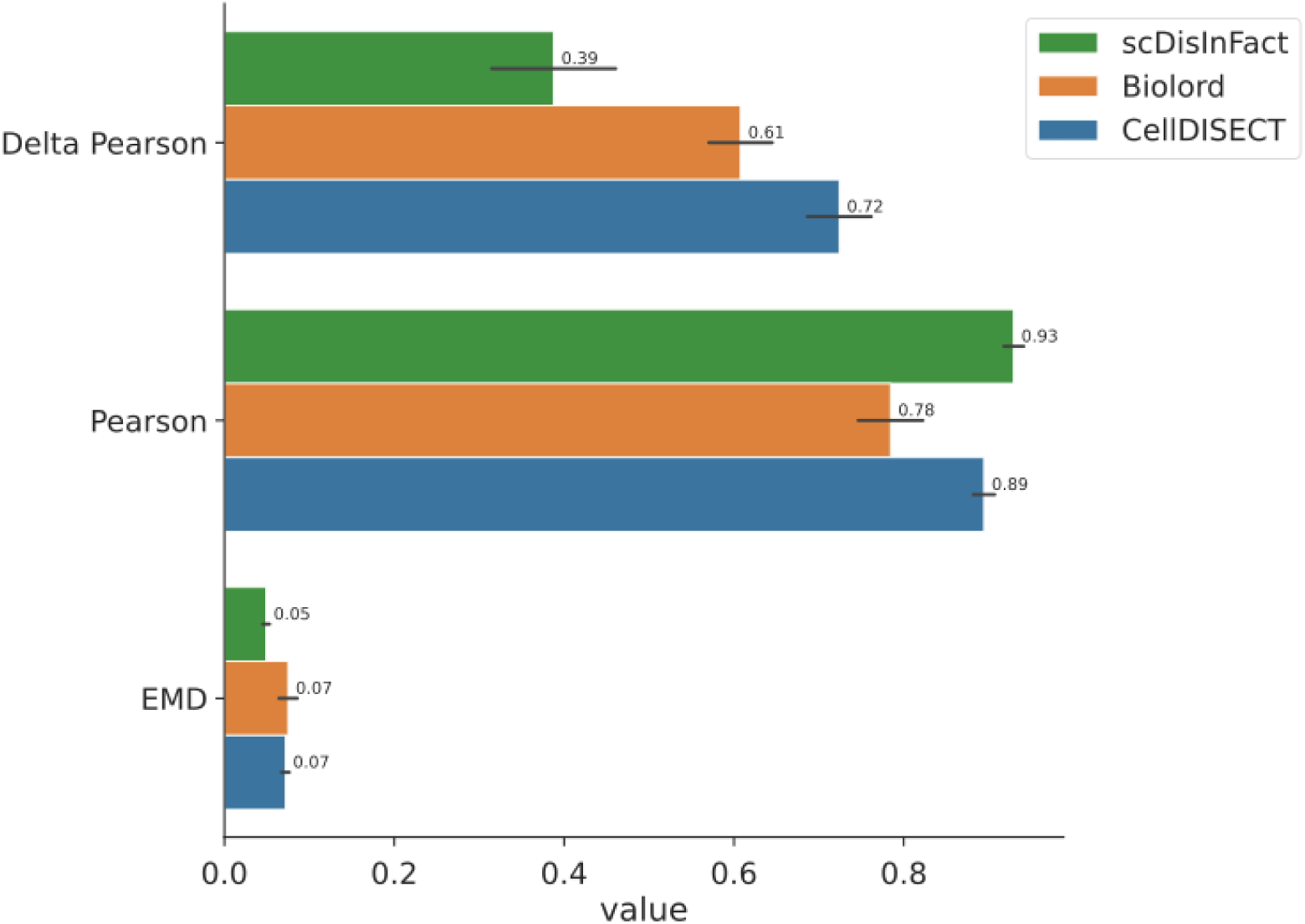
Kang all genes. Benchmarking on the Blood dataset of three models (scDisInFact, Biolord, and CellDISECT) on in-vivo vs. in-silico perturbation prediction across three evaluation metrics: Delta Pearson, Pearson, and EMD. Bars represent performance scores, with higher Pearson and Delta Pearson values indicating better agreement between in-vivo and in-silico perturbations. The Earth Mover’s Distance (EMD) reflects distributional similarity, where lower values indicate better performance. Error bars denote standard deviations. The reported Pearson correlation is between in-silico perturbed and in-vivo perturbed cells across all genes. The Delta Pearson reported is the correlation between in-silico perturbed and in-vivo perturbed after the unperturbed gene expression profile has been subtracted from both. The Earth Mover’s Distance (EMD) is the average of the dissimilarities between in-silico perturbed and in-vivo perturbed cells in gene space.

**Supplementary figure 2.**
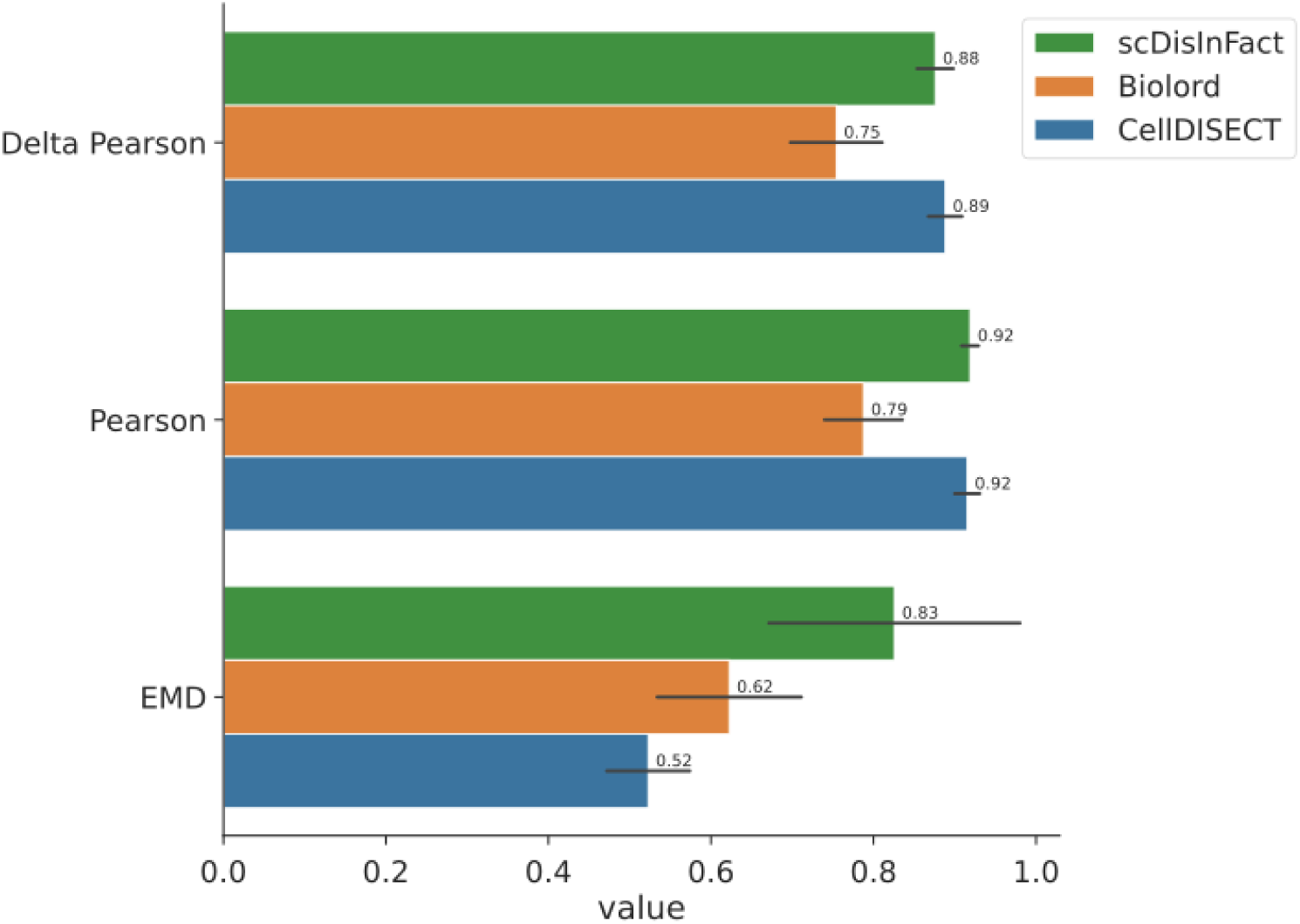
Kang 20 DEGs. Benchmarking on the Blood dataset of three models (scDisInFact, Biolord, and CellDISECT) on in-vivo vs. in-silico perturbation prediction across three evaluation metrics: Delta Pearson, Pearson, and EMD. Bars represent performance scores, with higher Pearson and Delta Pearson values indicating better agreement between in-vivo and in-silico perturbations. The Earth Mover’s Distance (EMD) reflects distributional similarity, where lower values indicate better performance. Error bars denote standard deviations. The reported Pearson correlation is between in-silico perturbed and in-vivo perturbed cells across the top 20 differentially expressed (between control and in-vivo perturbed) genes. The Delta Pearson reported is the correlation between in-silico perturbed and in-vivo perturbed after the unperturbed gene expression profile has been subtracted from both. The Earth Mover’s Distance (EMD) is the average of the dissimilarities between in-silico perturbed and in-vivo perturbed cells in gene space.

**Supplementary figure 3.**
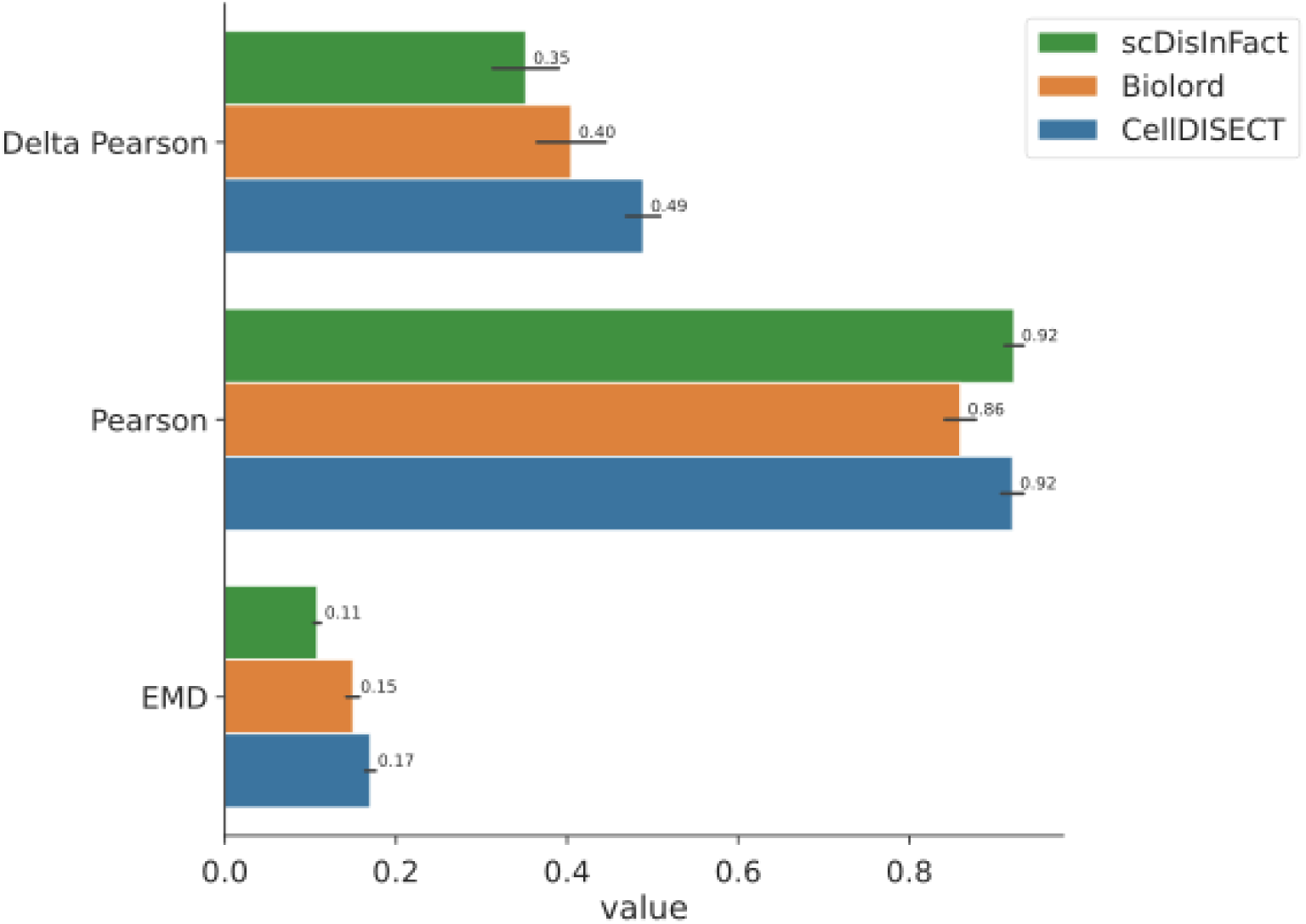
Haber all genes. Benchmarking on the Haber dataset of three models (scDisInFact, Biolord, and CellDISECT) on in-vivo vs. in-silico perturbation prediction across three evaluation metrics: Delta Pearson, Pearson, and EMD. Bars represent performance scores, with higher Pearson and Delta Pearson values indicating better agreement between in-vivo and in-silico perturbations. The Earth Mover’s Distance (EMD) reflects distributional similarity, where lower values indicate better performance. Error bars denote standard deviations. The reported Pearson correlation is between in-silico perturbed and in-vivo perturbed cells across all genes. The Delta Pearson reported is the correlation between in-silico perturbed and in-vivo perturbed after the unperturbed gene expression profile has been subtracted from both. The Earth Mover’s Distance (EMD) is the average of the dissimilarities between in-silico perturbed and in-vivo perturbed cells in gene space.

**Supplementary figure 4.**
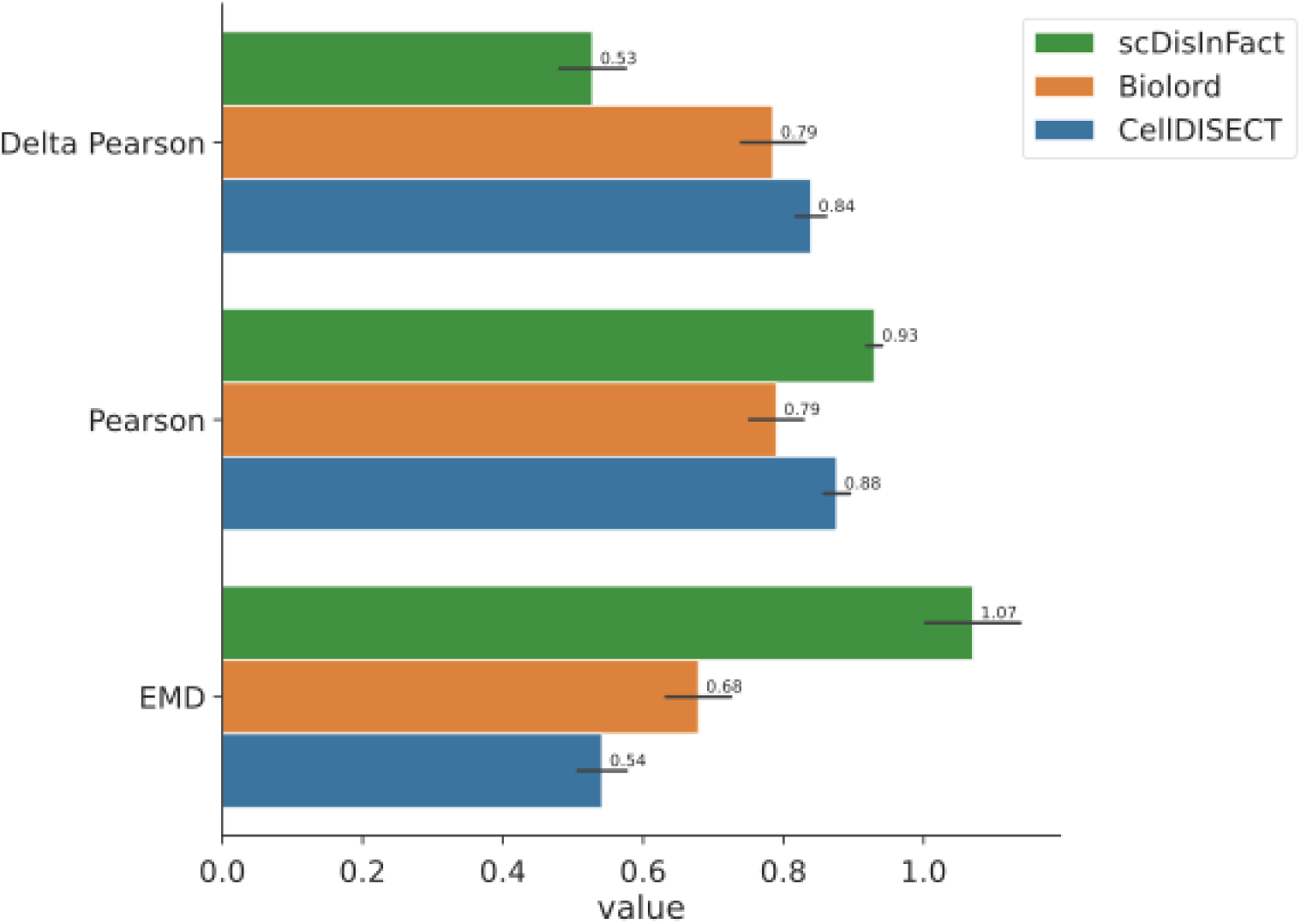
Haber 20 DEGs. Benchmarking on the Haber dataset of three models (scDisInFact, Biolord, and CellDISECT) on in-vivo vs. in-silico perturbation prediction across three evaluation metrics: Delta Pearson, Pearson, and EMD. Bars represent performance scores, with higher Pearson and Delta Pearson values indicating better agreement between in-vivo and in-silico perturbations. The Earth Mover’s Distance (EMD) reflects distributional similarity, where lower values indicate better performance. Error bars denote standard deviations. The reported Pearson correlation is between in-silico perturbed and in-vivo perturbed cells across the top 20 differentially expressed (between control and in-vivo perturbed) genes. The Delta Pearson reported is the correlation between in-silico perturbed and in-vivo perturbed after the unperturbed gene expression profile has been subtracted from both. The Earth Mover’s Distance (EMD) is the average of the dissimilarities between in-silico perturbed and in-vivo perturbed cells in gene space.

**Supplementary figure 5.**
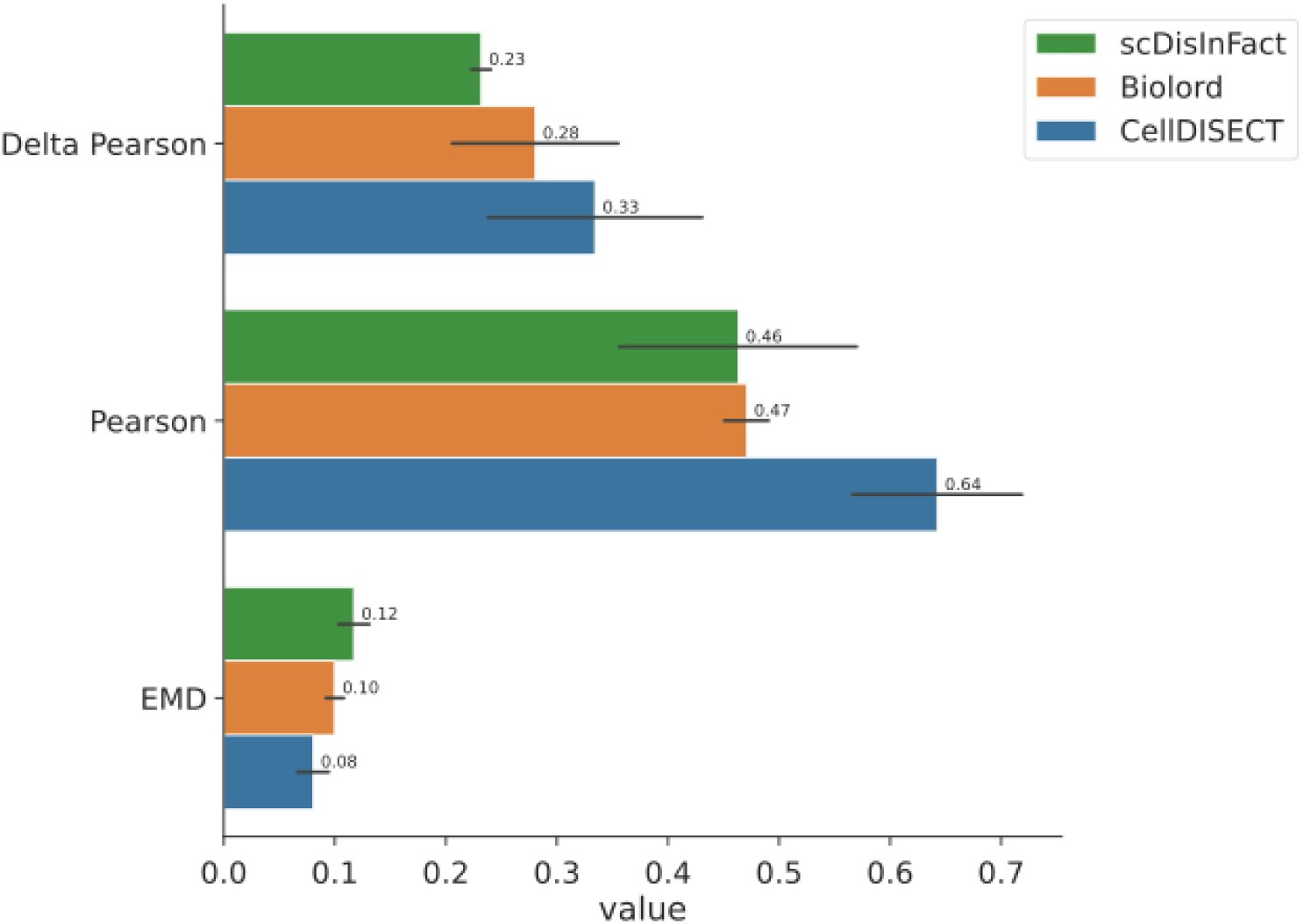
Eraslan all genes. Benchmarking on the Eraslan dataset of three models (scDisInFact, Biolord, and CellDISECT) on in-vivo vs. in-silico perturbation prediction across three evaluation metrics: Delta Pearson, Pearson, and EMD. Bars represent performance scores, with higher Pearson and Delta Pearson values indicating better agreement between in-vivo and in-silico perturbations. The Earth Mover’s Distance (EMD) reflects distributional similarity, where lower values indicate better performance. Error bars denote standard deviations. The reported Pearson correlation is between in-silico perturbed and in-vivo perturbed cells across all genes. The Delta Pearson reported is the correlation between in-silico perturbed and in-vivo perturbed after the unperturbed gene expression profile has been subtracted from both. The Earth Mover’s Distance (EMD) is the average of the dissimilarities between in-silico perturbed and in-vivo perturbed cells in gene space.

**Supplementary figure 6.**
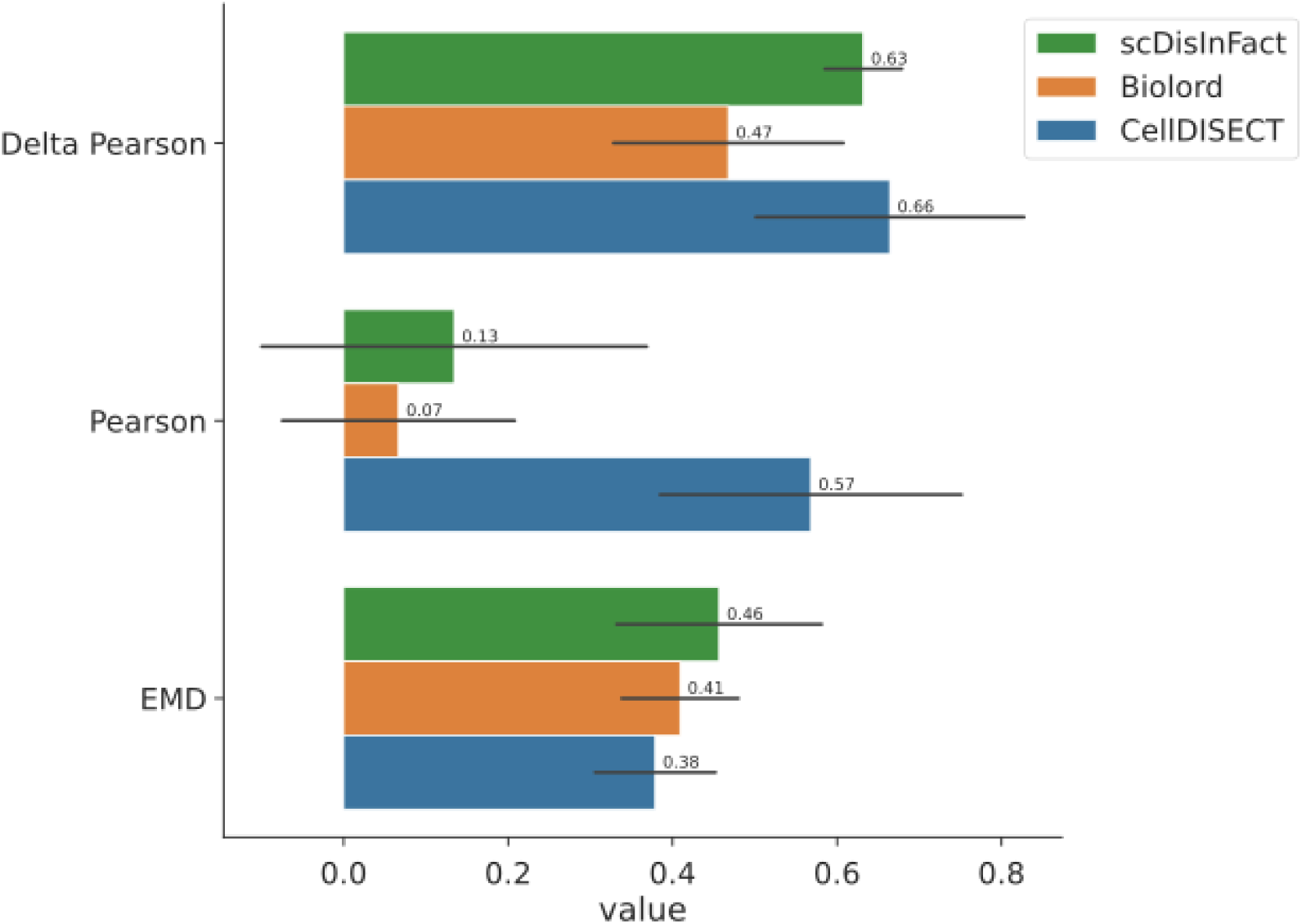
Eraslan top20 DEGs. Benchmarking on the Eraslan dataset of three models (scDisInFact, Biolord, and CellDISECT) on in-vivo vs. in-silico perturbation prediction across three evaluation metrics: Delta Pearson, Pearson, and EMD. The reported values are averages across all perturbation scenarios in the Eraslan dataset.. Bars represent performance scores, with higher Pearson and Delta Pearson values indicating better agreement between in-vivo and in-silico perturbations. The Earth Mover’s Distance (EMD) reflects distributional similarity, where lower values indicate better performance. Error bars denote standard deviations. The reported Pearson correlation is between in-silico perturbed and in-vivo perturbed cells across the top 20 differentially expressed (between control and in-vivo perturbed) genes. The Delta Pearson reported is the correlation between in-silico perturbed and in-vivo perturbed after the unperturbed gene expression profile has been subtracted from both. The Earth Mover’s Distance (EMD) is the average of the dissimilarities between in-silico perturbed and in-vivo perturbed cells in gene space.

**Supplementary figure 7.**
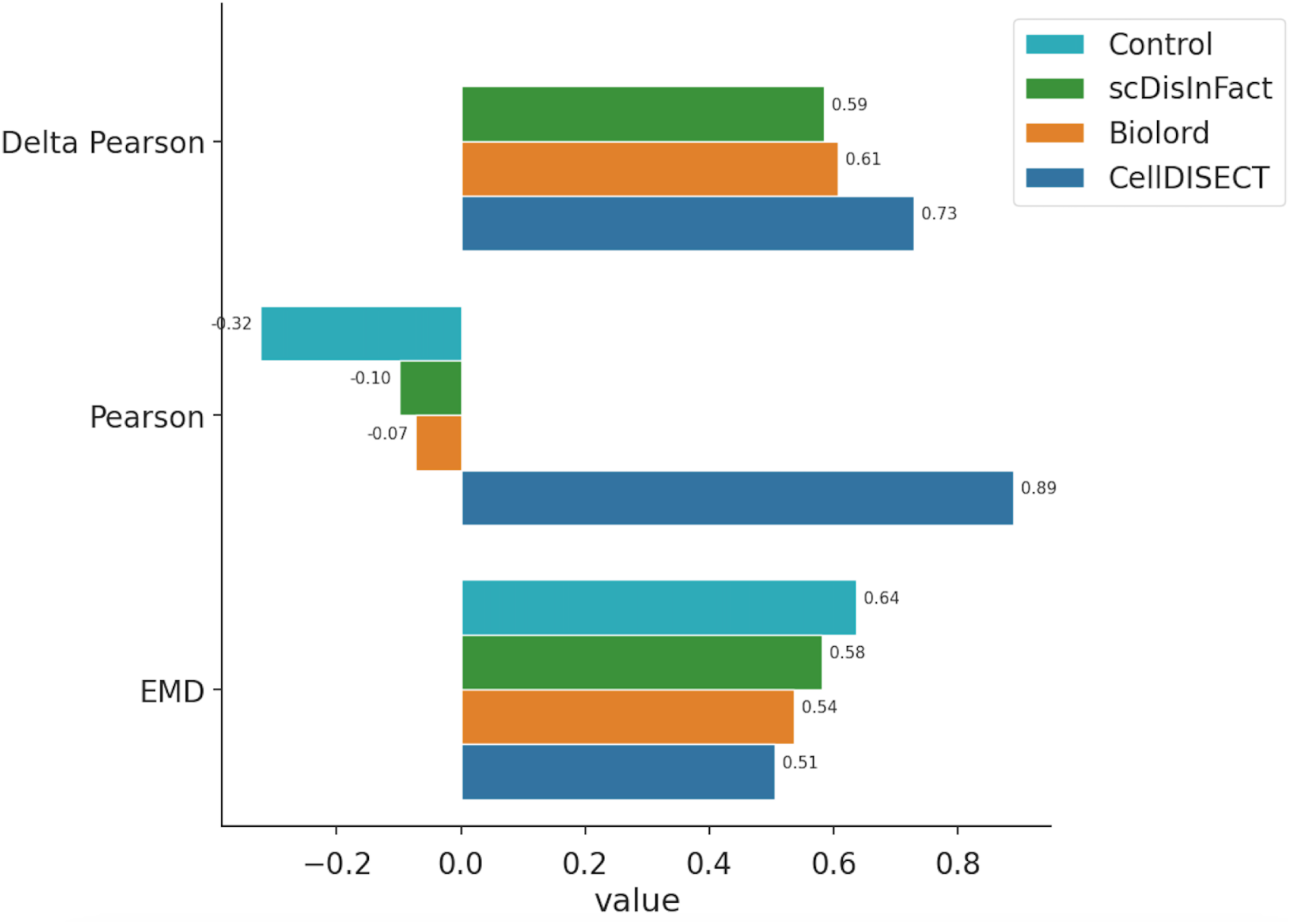
Eraslan scenario 2, female breast to male prostate. Benchmarking on the second perturbation scenario in the Eralsan dataset of three models (scDisInFact, Biolord, and CellDISECT) on in-vivo vs. in-silico perturbation prediction across three evaluation metrics: Delta Pearson, Pearson, and EMD. The perturbation task was a double counterfactual from epithelial cells in the female breast to male epithelial cells in the prostate. Bars represent performance scores, with higher Pearson and Delta Pearson values indicating better agreement between in-vivo and in-silico perturbations. The Earth Mover’s Distance (EMD) reflects distributional similarity, where lower values indicate better performance. Error bars denote standard deviations. The reported Pearson correlation is between in-silico perturbed and in-vivo perturbed cells across the top 20 differentially expressed (between control and in-vivo perturbed) genes. The Delta Pearson reported is the correlation between in-silico perturbed and in-vivo perturbed after the unperturbed gene expression profile has been subtracted from both. The Earth Mover’s Distance (EMD) is the average of the dissimilarities between in-silico perturbed and in-vivo perturbed cells in gene space.

**Supplementary figure 8.**
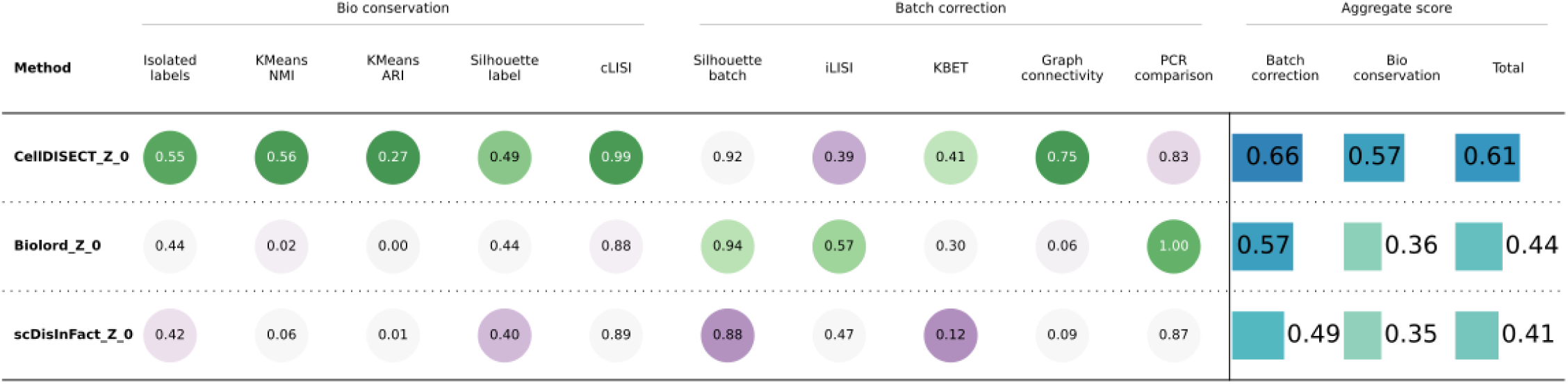
Benchmarking of the Z_0_ representation of the Eraslan^23^ dataset by three models (CellDISECT, Biolord, scDisInFact) using scIB metrics for evaluating single-cell data integration. The metrics are grouped into bio conservation, batch correction, and aggregate scores. Bio conservation assesses how well biological signals are preserved post-integration; batch correction evaluates the removal of batch effects; and aggregate scores summarize performance across bio conservation and batch correction, with higher scores indicating better overall integration quality.

**Supplementary figure 9.**
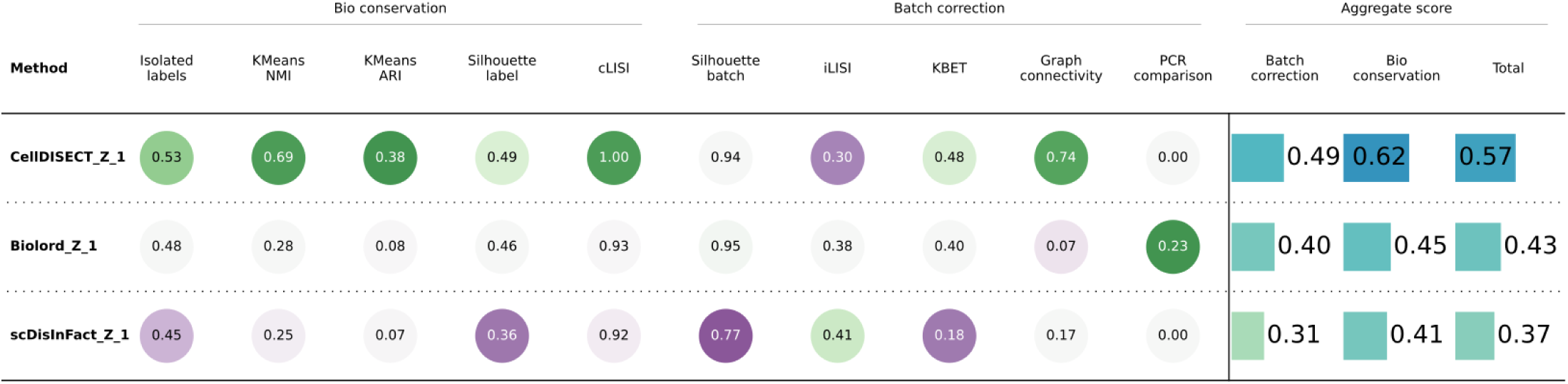
Benchmarking of the Z_tissue_ representation of the Eraslan ^23^ dataset by three models (CellDISECT, Biolord, scDisInFact) using scIB metrics for evaluating single-cell data integration. The metrics are grouped into bio conservation, batch correction, and aggregate scores. Bio conservation assesses how well biological signals are preserved post-integration; batch correction evaluates the removal of batch effects; and aggregate scores summarize performance across bio conservation and batch correction, with higher scores indicating better overall integration quality.

**Supplementary figure 10.**
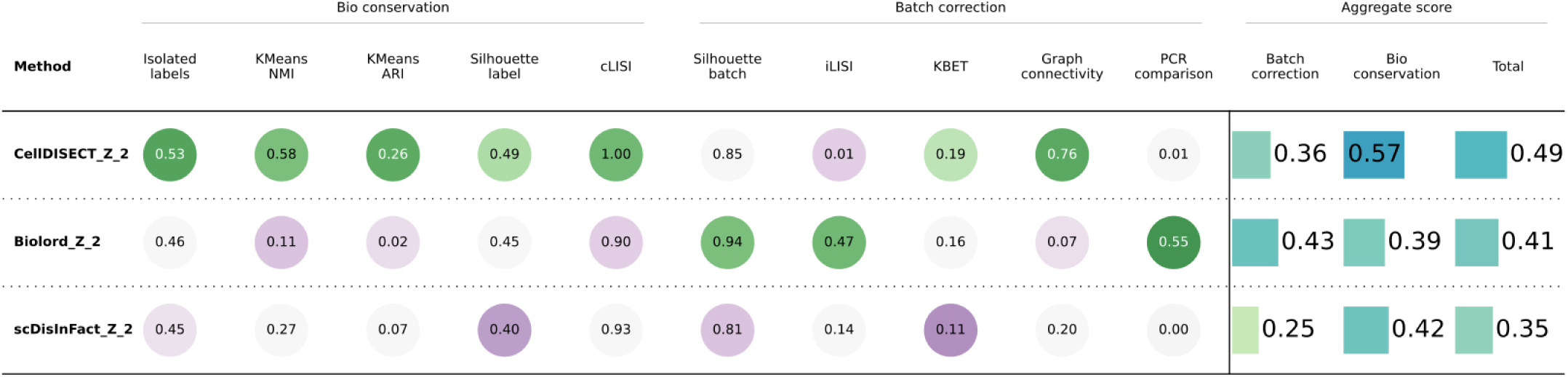
Benchmarking of the Z_sample ID_ representation of the Eraslan^23^ dataset by three models (CellDISECT, Biolord, scDisInFact) using scIB metrics for evaluating single-cell data integration. The metrics are grouped into bio conservation, batch correction, and aggregate scores. Bio conservation assesses how well biological signals are preserved post-integration; batch correction evaluates the removal of batch effects; and aggregate scores summarize performance across bio conservation and batch correction, with higher scores indicating better overall integration quality.

**Supplementary figure 11.**
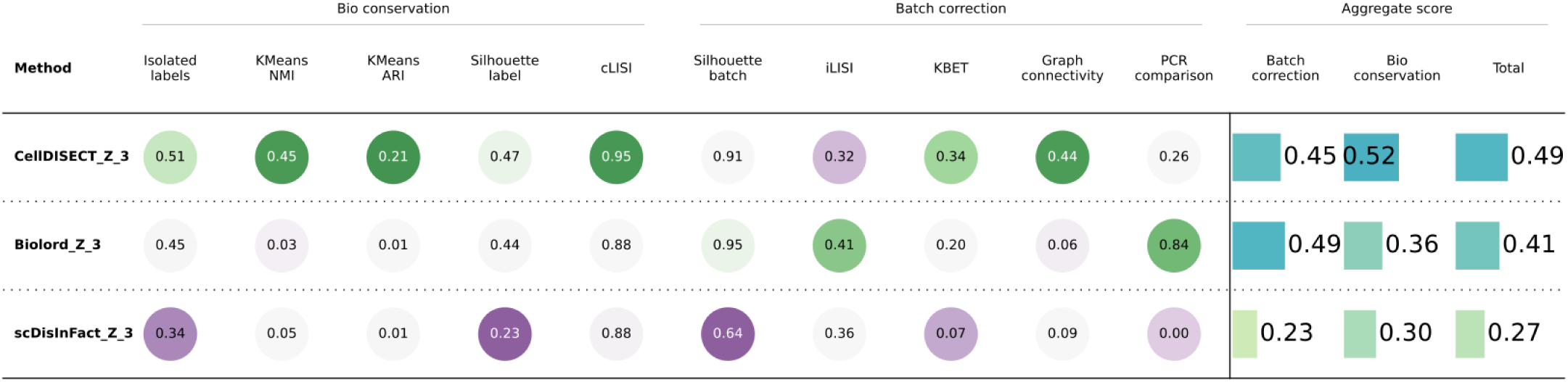
Benchmarking of the Z_sex_ representation of the Eraslan^23^ dataset by three models (CellDISECT, Biolord, scDisInFact) using scIB metrics for evaluating single-cell data integration. The metrics are grouped into bio conservation, batch correction, and aggregate scores. Bio conservation assesses how well biological signals are preserved post-integration; batch correction evaluates the removal of batch effects; and aggregate scores summarize performance across bio conservation and batch correction, with higher scores indicating better overall integration quality.

**Supplementary figure 12.**
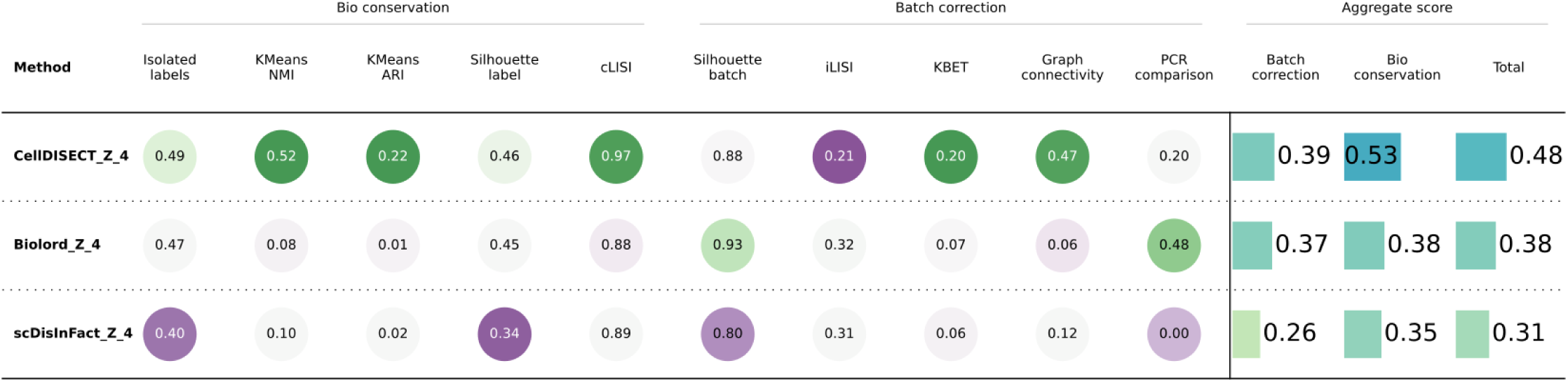
Benchmarking of the Z_age_ representation of the Eraslan^23^ dataset by three models (CellDISECT, Biolord, scDisInFact) using scIB metrics for evaluating single-cell data integration. The metrics are grouped into bio conservation, batch correction, and aggregate scores. Bio conservation assesses how well biological signals are preserved post-integration; batch correction evaluates the removal of batch effects; and aggregate scores summarize performance across bio conservation and batch correction, with higher scores indicating better overall integration quality.

**Supplementary figure 13.**
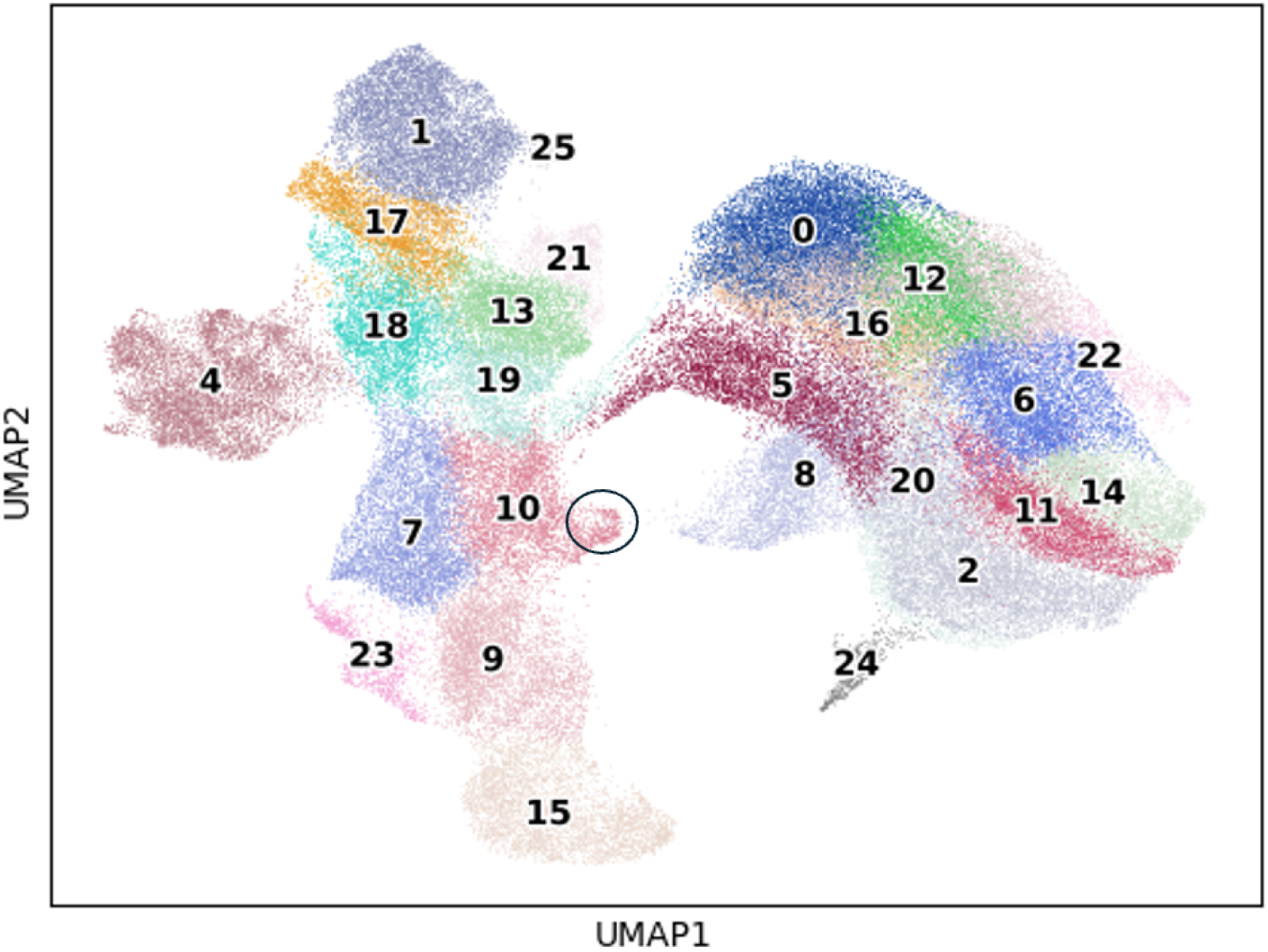
UMAP visualization of MEM lineage from the fetal immune atlas by Suo et al. colored by Leiden clustering at (default) resolution=1. The SPINK4+ subpopulation is denoted with a circle on the UMAP. Scvi does not identify the non-classical MK population which might be one contributing factor why this subpopulation was annotated as “early MK” by the original authors.

**Supplementary figure 14.**
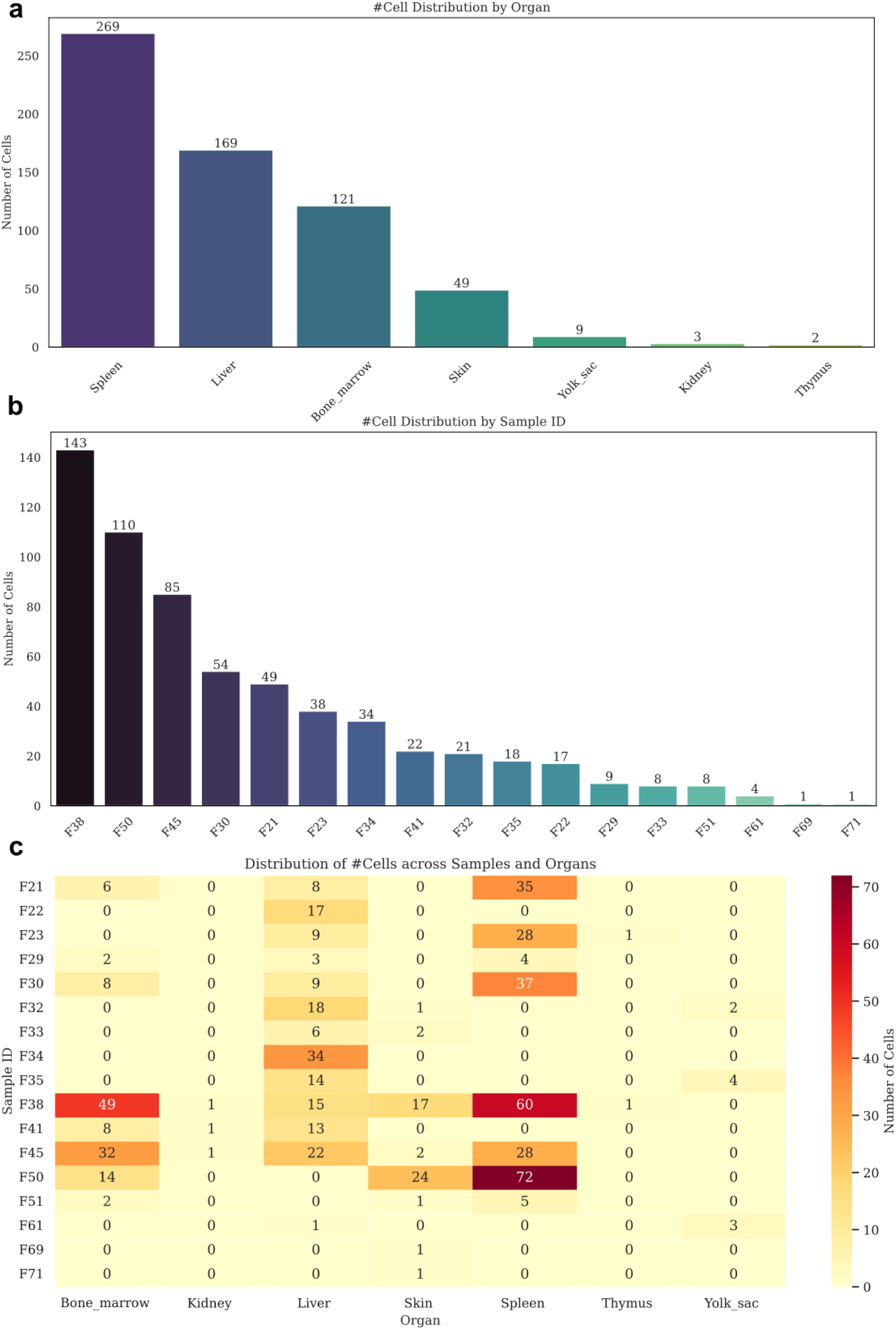
The novel SPINK4+ subpopulation is identified across several organs and sample IDs. a) Cell distribution by organ: Bar plot showing the number of SPINK4+ cells identified in various organs. The spleen contains the highest number of cells, followed by the liver and bone marrow, with fewer cells observed in the skin, yolk sac, kidney, and thymus. b) Cell distribution by sample ID: Bar plot displaying the number of SPINK4+ cells across individual samples, highlighting variability in cell counts among different biological samples. c) Heatmap of cell distribution across organs and sample IDs. Each row represents a sample ID, and each column represents an organ. The color intensity indicates the number of SPINK4+ cells per organ-sample combination, with darker shades representing higher cell counts.

**Supplementary figure 15.**
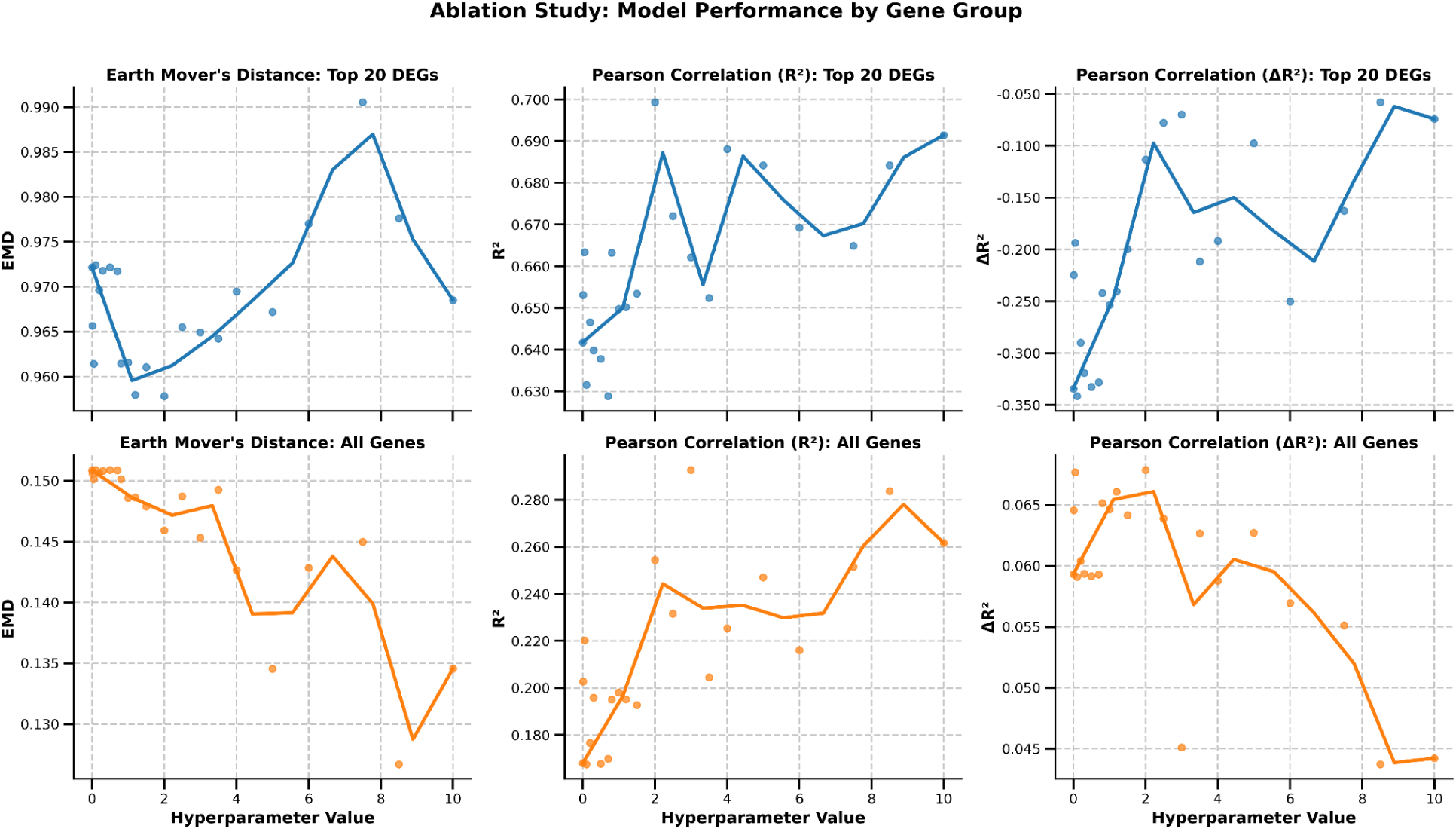
Ablation experiment of the effect of the counterfactual term on performance of counterfactual predictions at *test* time on the Blood dataset. Lines represent performance scores, with higher Pearson and Delta Pearson values indicating better agreement between in-vivo and in-silico perturbations. The Earth Mover’s Distance (EMD) reflects distributional similarity, where lower values indicate better performance. The reported Pearson correlation is between in-silico perturbed and in-vivo perturbed cells across all the highly variable genes, as well as the top 20 differentially expressed (between control and in-vivo perturbed) genes. The Delta Pearson reported is the correlation between in-silico perturbed and in-vivo perturbed after the unperturbed gene expression profile has been subtracted from both. The Earth Mover’s Distance (EMD) is the average of the dissimilarities between in-silico perturbed and in-vivo perturbed cells in gene space.

**Supplementary figure 16.**
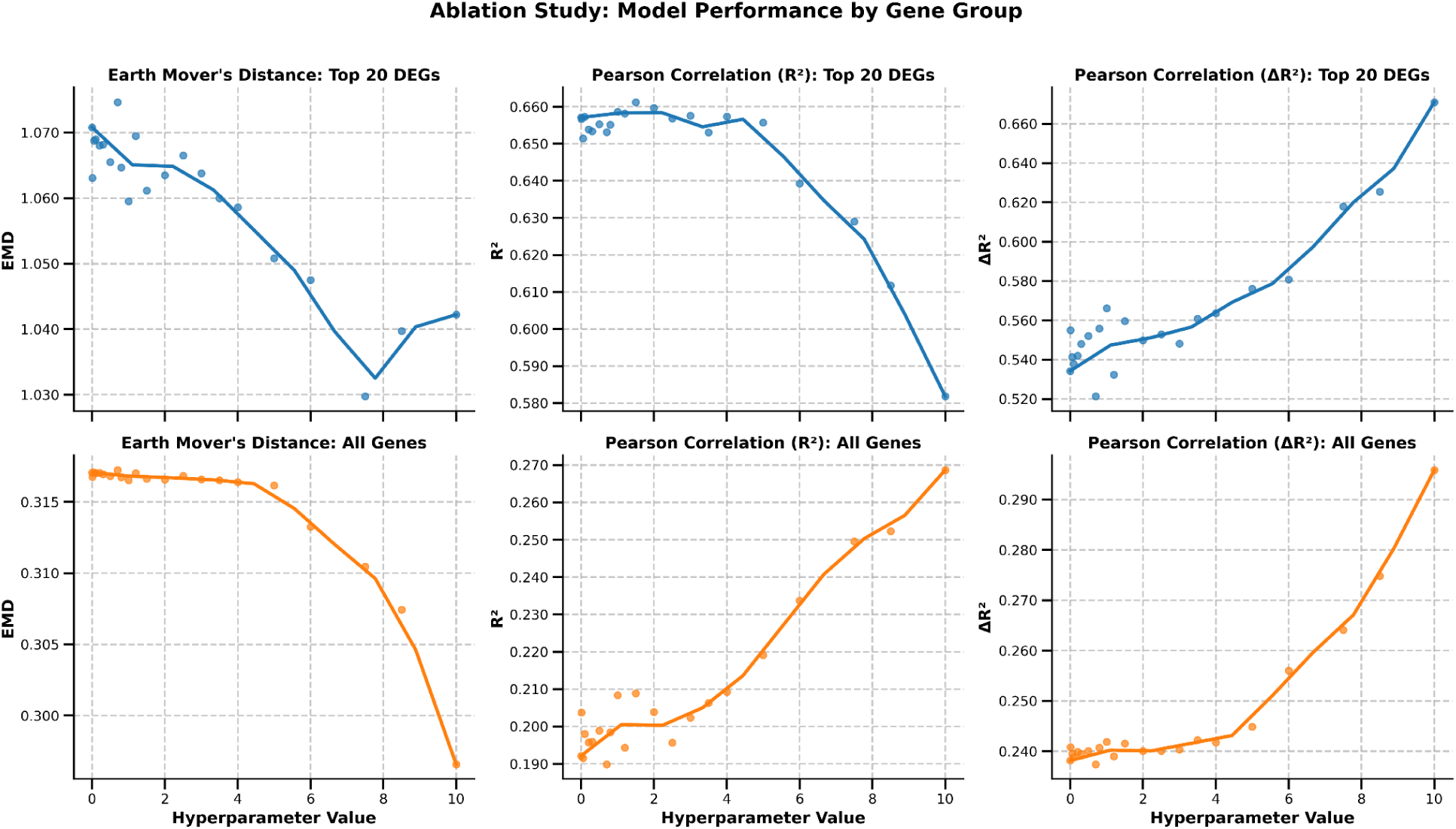
Ablation experiment of the effect of the counterfactual term on performance of counterfactual predictions at *test* time on the Intestine dataset. Lines represent performance scores, with higher Pearson and Delta Pearson values indicating better agreement between in-vivo and in-silico perturbations. The Earth Mover’s Distance (EMD) reflects distributional similarity, where lower values indicate better performance. The reported Pearson correlation is between in-silico perturbed and in-vivo perturbed cells across all the highly variable genes, as well as the top 20 differentially expressed (between control and in-vivo perturbed) genes. The Delta Pearson reported is the correlation between in-silico perturbed and in-vivo perturbed after the unperturbed gene expression profile has been subtracted from both. The Earth Mover’s Distance (EMD) is the average of the dissimilarities between in-silico perturbed and in-vivo perturbed cells in gene space.

**Supplementary figure 17.**
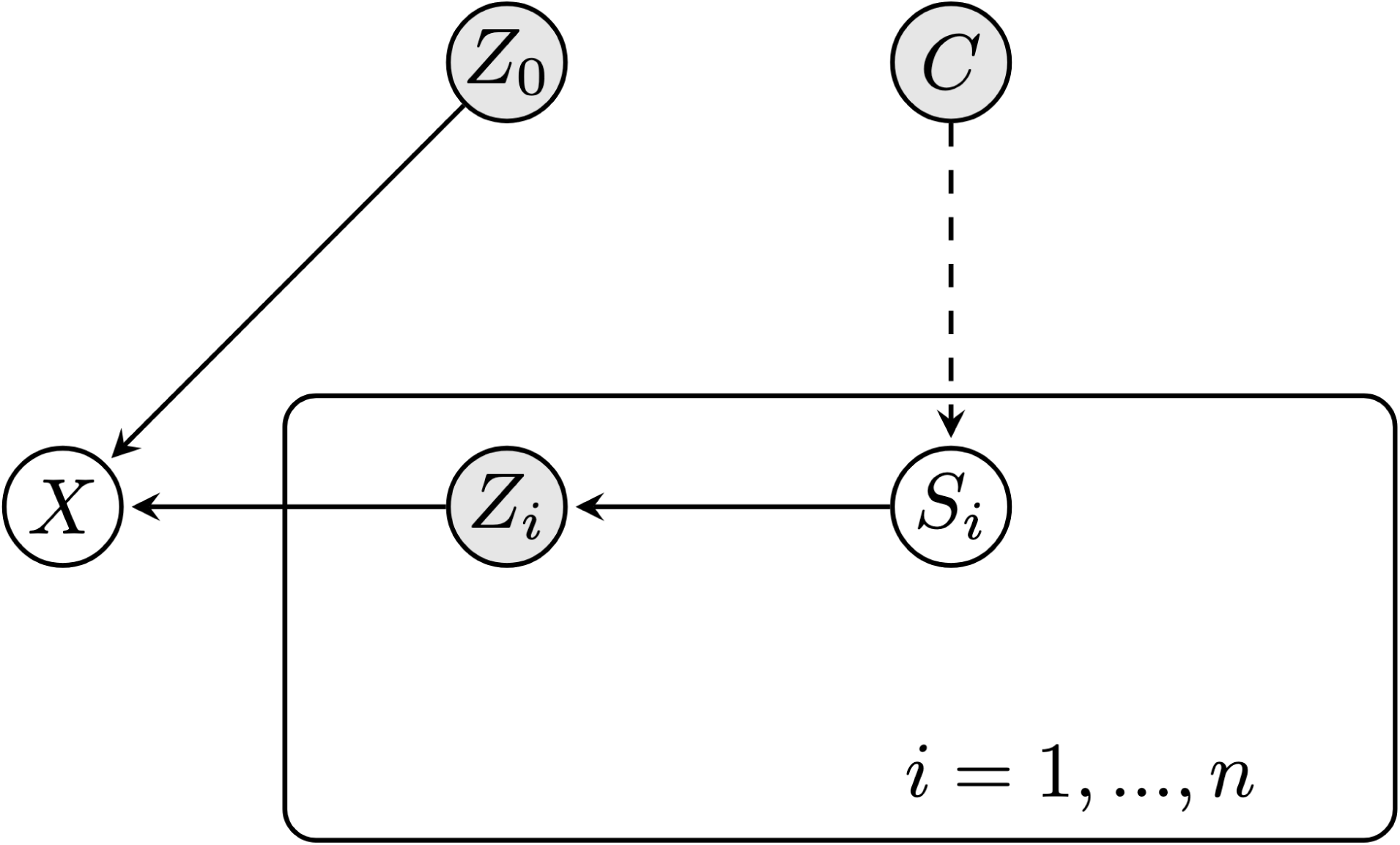
Structural Causal Model. Structural Causal Model for covariates Si and one response variable X with mediating latent variables Zi, where i = 1, …, n. Grey nodes are hidden, white nodes are observed, and C denotes potential unobserved confounders of the covariates.

## References

1. Klein, A. M. et al. Droplet barcoding for single-cell transcriptomics applied to embryonic stem cells. Cell 161, 1187–1201 (2015).

2. Macosko, E. Z. et al. Highly Parallel Genome-wide Expression Profiling of Individual Cells Using Nanoliter Droplets. Cell 161, 1202–1214 (2015).

3. Lotfollahi, M. et al. Predicting cellular responses to complex perturbations in high-throughput screens. Mol. Syst. Biol. 19, e11517 (2023).

4. Wang, X., Chen, H., Tang, S. ’ao, Wu, Z. & Zhu, W. Disentangled Representation Learning. IEEE Trans. Pattern Anal. Mach. Intell. 46, 9677–9696 (2024).

5. Chen, R. T. Q., Li, X., Grosse, R. & Duvenaud, D. Isolating sources of disentanglement in variational autoencoders. arXiv [cs.LG] (2018).

6. Pearl, J. Causality. (Cambridge University Press, 2009).

7. Feuerriegel, S. et al. Causal machine learning for predicting treatment outcomes. Nat. Med. 30, 958–968 (2024).

8. Liu, R., et al. Best practices and lessons learned on synthetic data for language models. ArXiv abs/2404.07503, (2024).

9. van Breugel, B. & van der Schaar, M. Beyond privacy: Navigating the opportunities and challenges of synthetic data. arXiv [cs.LG] (2023).

10. DeepSeek-AI, et al. DeepSeek-R1: Incentivizing Reasoning Capability in LLMs via Reinforcement Learning. arXiv [cs.CL] (2025).

11. Shumailov, I. et al. AI models collapse when trained on recursively generated data. Nature 631, 755–759 (2024).

12. Dohmatob, E., Feng, Y., Subramonian, A. & Kempe, J. Strong model collapse. arXiv [cs.LG] (2024).

13. Piran, Z., Cohen, N., Hoshen, Y. & Nitzan, M. Disentanglement of single-cell data with biolord. Nat Biotechnol 42, 1678–1683 (2024).

14. Zhang, Z., Zhao, X., Bindra, M., Qiu, P. & Zhang, X. scDisInFact: disentangled learning for integration and prediction of multi-batch multi-condition single-cell RNA-sequencing data. Nat Commun 15, 912 (2024).

15. Foster, A. et al. Contrastive Mixture of posteriors for counterfactual inference, data integration and fairness. arXiv [stat.ML] (2021).

16. Wu, Y., et al. Variational Causal Inference. arXiv [stat.ML] (2022).

17. Weinberger, E., Lin, C. & Lee, S.-I. Isolating salient variations of interest in single-cell data with contrastiveVI. Nat Methods 20, 1336–1345 (2023).

18. Suo, C. et al. Mapping the developing human immune system across organs. Science (2022) doi:10.1126/science.abo0510.

19. Wang, H. et al. Decoding human megakaryocyte development. Cell Stem Cell 28, 535–549.e8 (2021).

20. Kingma, D. P. & Welling, M. An Introduction to Variational Autoencoders. (2019).

21. Creager, E. et al. Flexibly Fair Representation Learning by Disentanglement. in International Conference on Machine Learning 1436–1445 (PMLR, 2019).

22. Website. https://github.com/Lotfollahi-lab/CellDISECT.

23. Eraslan, G. et al. Single-nucleus cross-tissue molecular reference maps toward understanding disease gene function. Science 376, eabl4290 (2022).

24. Luecken, M. D. et al. Benchmarking atlas-level data integration in single-cell genomics. Nat. Methods 19, 41–50 (2022).

25. Kang, H. M. et al. Multiplexed droplet single-cell RNA-sequencing using natural genetic variation. Nature Biotechnology 36, 89–94 (2017).

26. Haber, A. L. et al. A single-cell survey of the small intestinal epithelium. Nature 551, 333–339 (2017).

27. Heumos, L. et al. Best practices for single-cell analysis across modalities. Nat. Rev. Genet. 24, 550–572 (2023).

28. Cui, H. et al. scGPT: toward building a foundation model for single-cell multi-omics using generative AI. Nat. Methods 21, 1470–1480 (2024).

29. Vitrat, N. et al. Endomitosis of human megakaryocytes are due to abortive mitosis. Blood 91, (1998).

30. Liu, C. et al. Characterization of Cellular Heterogeneity and an Immune Subpopulation of Human Megakaryocytes. Advanced Science 8, 2100921 (2021).

31. Pariser, D. N. et al. Lung megakaryocytes are immune modulatory cells. The Journal of Clinical Investigation 131, e137377 (2021).

32. Hart, A. et al. Fli-1 is required for murine vascular and megakaryocytic development and is hemizygously deleted in patients with thrombocytopenia. Immunity 13, 167–177 (2000).

33. Doré, L. C. & Crispino, J. D. Transcription factor networks in erythroid cell and megakaryocyte development. Blood 118, 231–239 (2011).

34. Psaila, B. et al. Single-Cell Analyses Reveal Megakaryocyte-Biased Hematopoiesis in Myelofibrosis and Identify Mutant Clone-Specific Targets. Mol Cell 78, 477–492.e8 (2020).

35. Bühring, H. J. et al. The basophil activation marker defined by antibody 97A6 is identical to the ectonucleotide pyrophosphatase/phosphodiesterase 3. Blood 97, 3303–3305 (2001).

36. Tsai, S. H. et al. The ectoenzyme E-NPP3 negatively regulates ATP-dependent chronic allergic responses by basophils and mast cells. Immunity 42, 279–293 (2015).

37. Krystel-Whittemore, M., Dileepan, K. N. & Wood, J. G. Mast Cell: A Multi-Functional Master Cell. Front Immunol 6, 620 (2015).

38. Anbazhagan, M. et al. PTGER4 signaling regulates class IIa HDAC function and SPINK4 mRNA levels in rectal epithelial cells. Cell Commun. Signal. 22, 493 (2024).

39. Hu, B.-L., Yin, Y.-X., Li, K.-Z., Li, S.-Q. & Li, Z. SPINK4 promotes colorectal cancer cell proliferation and inhibits ferroptosis. BMC Gastroenterol. 23, 104 (2023).

40. Wang, Y. et al. Therapeutic potential of the secreted Kazal-type serine protease inhibitor SPINK4 in colitis. Nat. Commun. 15, 5874 (2024).

41. Wells, M. & Steiner, L. Epigenetic and Transcriptional Control of Erythropoiesis. Front Genet 13, 805265 (2022).

42. Zhao, B., Yang, J. & Ji, P. Chromatin condensation during terminal erythropoiesis. Nucleus 7, 425–429 (2016).

43. Wong, P. et al. Gene induction and repression during terminal erythropoiesis are mediated by distinct epigenetic changes. Blood 118, e128–38 (2011).

44. Cellxgene Data Portal. Cellxgene Data Portal https://cellxgene.cziscience.com/.

45. Heumos, L., et al. Pertpy: an end-to-end framework for perturbation analysis. bioRxiv (2024) doi:10.1101/2024.08.04.606516.

46. Website. https://www.celltypist.org/encyclopedia/Human_Whole_Embryo/v1.

47. Wolf, F. A., Angerer, P. & Theis, F. J. SCANPY: large-scale single-cell gene expression data analysis. Genome Biology 19, 1–5 (2018).

48. Cellxgene Data Portal. Cellxgene Data Portal https://cellxgene.cziscience.com/.

49. Khemakhem, I., Kingma, D. P., Monti, R. P. & Hyvärinen, A. Variational Autoencoders and Nonlinear ICA: A Unifying Framework. arXiv [stat.ML] (2019).

