## Supplementary Note 1 for "Integrating multi-covariate disentanglement with counterfactual analysis on synthetic data enables cell type discovery and counterfactual predictions"

### SUPPLEMENTARY NOTE 1: THEORETICAL FOUNDATIONS OF DISENTANGLED EXPERTS FOR COVARIATE COUNTERFACTUALS (CELLDISECT)

#### 1 THEORETICAL FOUNDATIONS OF CELLDISECT

CellDISECT estimates for cell $n$ the conditional distribution, $p(x_n|s_{1n}, ..., s_{Cn})$ in which $x_n$ is the G-dimensional vector of observed RNA counts (G is the total number of genes), and $s_{1n}, ..., s_{Cn}$ are covariate vectors describing information such as batch index of cell, age of donor, tissue, etc. In total there are $N$ cells and $C$ covariates for each cell. This distribution is estimated using a Mixture of $C+1$ Expert (MoE) VAEs, and $C(C+1)$ MLPs that (de)couple these expert VAEs. In addition to interpretability, and flexible batch correction, this decoupling also offers a straightforward approach to counterfactual prediction as explained in Foster et al. (2022), (and we leverage this by training our algorithm to maximize the sum of the likelihood of the observed data and the likelihood of the model's counterfactual predictions).

Throughout the rest of the paper, when we refer to a specific cell or generally the random variable, we will drop the superscript $n$.

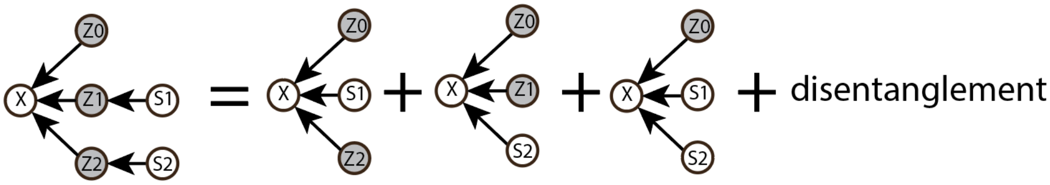


Figure 1: A cartoon depicting our approach to modelling single-cell data with many covariates and batches. By marginalizing $z_1$, then $z_2$, then both, we get reduced structural causal models each of which can be modeled with a conditional VAE. We call these VAEs expert VAEs because each is aligned with a specific covariate of interest.

##### 1.1 MOTIVATION

Our central idea is to provide users with a multifaceted view of the data with many different latent spaces and VAEs, each aligned with a different covariate.

We get each of these expert VAEs by mathematically marginalizing out all covariates to which we don't want to align, as shown in the cartoon of Fig 1.

##### 1.2 MODEL DERIVATION

Our model of covariate-specific experts naturally follows from our assumed structural causal model (SCM) generating our data (see Fig. 2), and Proposition 1.

**Proposition 1.** *The factorization of the SCM in figure 2 implies the following evidence lower bound,*

$$\log p(x|s) \geq \frac{1}{C+1} \sum_{i=0}^{C} \left[ \mathbb{E}_{q_{\phi_i}(z_i|s,x)}[\log p(z_i|s_i) - \log q_{\phi_i}(z_i|s,x)] + \mathbb{E}_{q_{\phi_i}(z_i|s,x)}[\log p(x|z_i, s_{-i}] \right], \tag{1}$$

*where $s = \cup_i s_i$, $s_{-i} = s - \{s_i\}$, and $q_{\phi_i}$ is any encoding function.*

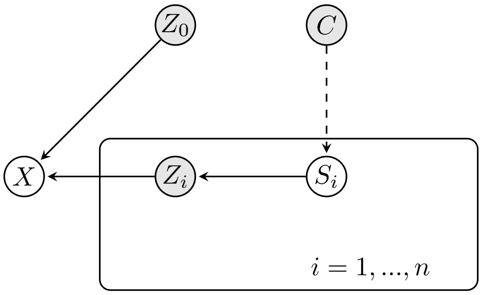


Figure 2: (Left) Structural Causal Model for covariates $S_i$ and one response variable X with mediating latent variables $Z_i$, where $i = 1, ..., n$. Grey nodes are hidden, white nodes are observed, and C denotes potential confounders of the covariates.

*Proof.*

$$\begin{aligned}
\log p(x|s) &= \mathbb{E}_{q_{\phi_i}(z_i|s,x)}[\log p(x|s)] && (2)\\
&= \mathbb{E}_{q_{\phi_i}(z_i|s,x)}[\log p(x, z_i|s) - \log p(z_i|s,x)] && (3)\\
&= \mathbb{E}_{q_{\phi_i}(z_i|s,x)}[\log p(x, z_i|s) - \log p(z_i|s,x) + (q_{\phi_i}(z_i|s,x) - q_{\phi_i}(z_i|s,x))] && (4)\\
&= D(q_{\phi_i}(z_i|s,x) \parallel p(z_i|s,x)) + \mathbb{E}_{q_{\phi_i}(z_i|s,x)}\Big[\log p(x, z_i|s) - q_{\phi_i}(z_i|s,x))\Big] && (5)\\
&\geq \mathbb{E}_{q_{\phi_i}(z_i|s,x)}\Big[\log p(z_i|s) - q_{\phi_i}(z_i|s,x))\Big] + \mathbb{E}_{q_{\phi_i}(z_i|s,x)}\Big[\log p(x|z_i,s))\Big] && (6)
\end{aligned}$$

in the first line we multiply and divide by the function $q_{\phi_i}(z_i|s,x)$. Although at this stage we are free to choose any function, this will ultimately be the encoder of the i-th expert VAE. $q_{\phi_i}(z_i|s,x)$ is the most general choice of an encoder consistent with our assumed SCM since it includes all the observed variables, and it cannot be simplified based on d-separation in the SCM.

Moreover, by d-separation on the causality graph we have

$$\begin{aligned}
\forall i :& z_i \perp s_{-i}|s_i && (7)\\
\forall i :& x \perp s_i|z_i, s_{-i} && (8)
\end{aligned}$$

which simplify the derived ELBO into

$$\log p(x|s) \geq \mathbb{E}_{q_{\phi_i}(z_i|s,x)}\Big[\log p(z_i|s_i) - q_{\phi_i}(z_i|s,x))\Big] + \mathbb{E}_{q_{\phi_i}(z_i|s,x)}\Big[\log p(x|z_i,s_{-i}))\Big] \quad (9)$$

Summing over $i$ and rearranging yields the desired result. □

In the ELBO of proposition 1, there remain three (related by Bayes's Theorem) functions, $p(x|s), p(z_i|s_i), p(x|z_i, s_{-i})$, that we can learn using neural networks, which leads to the following parametrization

$$\log p_\nu(x|s) \geq \frac{1}{C+1}\sum_{i=0}^{C}\Big[\mathbb{E}_{q_\phi(z_i|s,x)}[\log p_{\chi_i}(z_i|s_i) - \log q_\phi(z_i|s,x)] + \mathbb{E}_{q_\phi(z_i|s,x)}[\log p_{\lambda_i}(x|z_i, s_{-i}]\Big], \quad (10)$$

where $\nu = \{\chi_i, \lambda_i : \forall i\}$. As proposition 1 shows, for our mixture of experts, we need $C+1$ encoders, $q_{\phi_i}(z_i|s,x)$, which take as input all the covariates and the gene expression; $C+1$ decoders, $p_{\lambda_i}(x|z_i, s_{-i})$, which take as input a latent variables, all the covariates but the one corrresponding the the latent variable; and we also use $C+1$ prior encoders, $p_{\chi_i}(z_i|s_i)$, to allow more flexibility in learning the prior distributions for the latent variables, which has been shown to ensure identifiability of the latent space variables in Khemakhem et al. (2020).

##### 1.3 DISENTANGLING THE COVARIATE-SPECIFIC EXPERTS

So far our model is a robust way to learn our desired conditional distribution, $p(x|s)$, but the latent spaces are not guaranteed to be causally disentangled Foster et al. (2022), in the sense that the learnt $z_{j\neq i}$ might be causally influenced by $s_i$. This is possible since the $i$-th encoder uses all of $s$ to calculate $z_i$. To causally disentangle the $z_i$ from the $s_{j\neq i}$, we follow a Granger causality mentality. In Granger (1969) he proposed a notion of causality based on how well past values of a time series $y_t$ can predict future values of another times series $x_t$. Granger defined y to be causal to x iff

$$\text{var}\big[x_t - f(x_t|x_{<t}, y_{<t})\big] < \text{var}\big[x_t - f(x_t|x_{<t})\big] \tag{11}$$

where $f(x|H)$ is the optimal predictor of $x$ given $H$. Due to the presence of confounders $C$ see figure 1, $s_i$ can be used to predict a $z_{j\neq i}$, however we think of $s_i$ as causally affecting a $z_{j\neq i}$ iff

$$CE(z_j, P(s_j, s_i)) < CE(z_j, P(s_j)). \tag{12}$$

where $CE(x, y)$ denotes the cross entropy between $x, y$ and $P$ is the optimal predictor of $z_j$.

To ensure that our covariate-specific representations $z_i$ are disentangled, which also makes counterfactual prediction more straightforward Foster et al. (2022), we introduce some auxiliary neural networks. This allows us to generalize the method in Foster et al. (2022) to cases with multiple conditions. More specifically we introduce $C$ classifiers (MLPs) that align $s_i$ to $z_i$ by learning to predict $s_i$ from $z_i$, and $C^2$ adversarial classifiers (MLPs) that erase information of $s_i$ in $z_{j\neq i}$ by learning to predict $s_i$ from $z_{j\neq i}$. As we show in Proposition 2, adding these auxiliary neural networks and their associated loss terms results in an objective function that still lower bounds $\log p(x|s)$. Moreover, as we show in proposition 3, these adversarial classifiers do not erase the information of $z_0$ which is contained in every $z_i$.

**Proposition 2.** *The sum of the (factual) posterior likelihood and the counterfactual posterior likelihood is lower bounded by*

$$\begin{aligned}
\log p(x|s) + \sum_{i=1}^{n} \log p(x'^{i}|x, s, s') \geq\ & \frac{1}{C+1} \sum_{i=0}^{C} \Big[ \mathbb{E}_{q_\phi(z_i|s,x)} [\log p_{\chi_i}(z_i|s_i) - \log q_\phi(z_i|s, x)] \Big] \\
& + \frac{1}{C+1} \sum_{i=0}^{C} \Big[ \mathbb{E}_{q_\phi(z_i|s,x)} [\log p_{\lambda_i}(x|z_i, s_{-i}] \Big] \\
& + \sum_{i=1}^{n} \log p(x'^{n}|x, s', s) \\
& - \sum_{i=1}^{n} CE(classifier(s_i|z_i), s_i) \\
& + \sum_{i=1}^{n} CE(classifier(s_i|z_{-i}), s_i) \\
& := -\mathcal{L}
\end{aligned} \tag{13}$$

*where $s = s_{-i} \cup s_i$, and $x'^{k}$ denotes a counterfactual with $k$ many interventions.*

*Proof.* The cross entropy of a any classifier of a categorical covariate $S_k$ with finite sample space of size $I$, is non-negative quantity with an upper bound determined by $I$,

$$0 \leq CE(\text{classifier}(s_i), s_i) \leq f(I). \tag{14}$$

This also holds for the cross entropy of continuous covariates whose distributions have compact support. Therefore from Proposition 1 and 14,

$$
\begin{aligned}
\log p_\nu(x|s) &\geq \frac{1}{C+1}\sum_{i=0}^{C}\Big[\mathbb{E}_{q_\phi(z_i|s,x)}[\log p_{\chi_i}(z_i|s_i) - \log q_\phi(z_i|s,x)] + \mathbb{E}_{q_\phi(z_i|s,x)}[\log p_{\lambda_i}(x|z_i,s_{-i}]\Big] \\
&\geq \frac{1}{C+1}\sum_{i=0}^{C}\Big[\mathbb{E}_{q_\phi(z_i|s,x)}[\log p_{\chi_i}(z_i|s_i) - \log q_\phi(z_i|s,x)] + \mathbb{E}_{q_\phi(z_i|s,x)}[\log p_{\lambda_i}(x|z_i,s_{-i}]\Big] \\
&- \sum_{i=1}^{n} CE(classifier(s_i|z_i), s_i) \\
&- \Big(f(I) - \sum_{i=1}^{n} CE(classifier(s_i|z_{-i}), s_i)\Big)
\end{aligned}
\tag{15}
$$

Now adding the term counterfactual likelihood for the n-intervention outcome $\sum_{i=1}^{n}\log p(x'|x,s,s')$ on both hand sides we obtain

$$
\log p(x|s) + \sum_{i=1}^{n}\log \hat{p}(x'^{n}|x,s',s) \geq -\mathcal{L} - f(I). \tag{16}
$$

Although the cross entropy depends on the classifier and its parameters its upper bound does not. It only depends on the number of possible values of the categorical covariate, and hence it is a constant that can be neglected in optimizing the parameter of our algorithm. This shows that minimizing the objective function in the main text $\mathcal{L}$ will increase the lower bound on the sum of the two log-likelihoods. □

##### 1.4 Counterfactual prediction and counterfactual semi-autoencoding

Having ensured that $z_j$ is causally independent from $s_i$, $\forall i \neq j$, counterfactual predictions are rendered more straightforward, as explained in Foster et al. (2022), by neatly separating the three steps in Pearl's method of counterfactual prediction Pearl (2009):

1) *abduction*: we infer the value of the root nodes $\{z_0\}$ based on the observations, $P(z_0|s,x)$,

2) *action*: swapping $s_i$ for $s'_i$, $do(s_i = s'_i)$,

3) *prediction*: use $P(x_{s'_i} = \tilde{x}|s'_i, s_i, s_{-i}, x)$ to obtain a prediction for $x^{cf}$ given the value for $z_0$ calculated in abduction.

The fundamental problem in causal inference is that we have access only to one outcome (termed factual). We never observe the counterfactual outcome, which is defined by the $do$ operations on the causality graph and therefore might not even be identifiable from observations. As we show in proposition 4, in our model, the counterfactual outcome of intervention on any covariate is identifiable by data that has been observed.

We now describe an alternative way of performing counterfactual predictions, which we empirically find to perform better, and use as the default version in our model. Intuitively, the approach below consists of making independent counterfactual predictions from all the decoders that are not associated with the perturbed covariates and then averaging the results. We now demonstrate that the counterfactuals can be identified by this approach as long as there are no unobserved confounders C (see Fig. 2).

**Proposition 3** (Identifiability2). *The counterfactual (never observed) value of the response variable $x$ due to an intervention $do(s_i = s'_i)$ on any covariate $i$ can be identified based solely on observed data and is equal to*

$$
P(x_{s'_i} = \tilde{x}|s'_i, s_i, s_{-i}, x) = \mathbb{E}_{p(z_j|s_i,s_{-i},s'_i,x)}\Big[P(x = \tilde{x}|s'_i, s_{-ij}, z_j)\Big], \tag{17}
$$

*where the RHS contains conditional probabilities that can be calculated from observed data.*

*Proof.*

$$P(x_{s_i'} = \tilde{x}|s_i', s_i, s_{-i}, x) = \int_{z_j} P(x_{s_i'} = \tilde{x}|s_i', s_i, s_{-i}, x, z_j) P(z_j|s_i, s_{-i}, s_i', x) dz_j \tag{18}$$

$$= \mathbb{E}_{p(z_j|s_i, s_{-i}, s_i', x)} \left[ P(x_{s_i'} = \tilde{x}|s_i', s_i, s_{-i}, x, z_j) \right] \tag{19}$$

$$= \mathbb{E}_{p(z_j|s_i, s_{-i}, s_i', x)} \left[ P(x_{s_i'} = \tilde{x}|s_i', s_i, s_{-ij}, z_j) \right] \tag{20}$$

$$= \mathbb{E}_{p(z_j|s_i, s_{-i}, s_i', x)} \left[ P(x_{s_i'} = \tilde{x}|s_i', s_{-ij}, z_j) \right] \tag{21}$$

$$= \mathbb{E}_{p(z_j|s_i, s_{-i}, s_i', x)} \left[ P(x = \tilde{x}|s_i', s_{-ij}, z_j) \right], \tag{22}$$

where in line 3 we used that

$$s_j \perp x_{s_i'}|z_j \tag{23}$$

$$x \perp x_{s_i'}|z_j, s_{-ij} \tag{24}$$

$$\tag{25}$$

Similarly, in line 4, we used that

$$s_i \perp x_{s_i'} \tag{26}$$

$$\tag{27}$$

These follow from d-separation on the parallel world graph 3. Finally, in line 5 we used the first property of counterfactuals (Composition). Pearl (2009). □

The quantity in the brackets, $P(x = \tilde{x}|s_i', s_{-ij}, z_j)$, is precisely the generative process of the $j$-th expert VAE, and $p(z_j|s_i, s_{-i}, s_i', x)$, is precisely the $j$-th encoder. This concludes our proof that the counterfactual expression $x_{s_i'}$ can be reduced to an ordinary probabilistic expression (involving no counterfactuals) learned by our method during training.

###### 1.4.1 SEMI-AUTOENCODING SYNTHETIC COUNTERFACTUALS

The identifiability of counterfactual predictions allows us to use the idea in Wu et al. (2022) of semi-autoencoding an algorithm's counterfactual predictions and jointly maximizing the likelihood of both real data and counterfactuals, $\log p(x|s) + \log p(x'|x, s, s')$, where $s' = s_{-i} \cup s_i'$. To implement the counterfactual semi-autoencoding we generalize Wu et al. (2022) to the case of multiple conditions.

Specifically, before training, we generate counterfactual pairs of cells. Every cell $c$ is paired with all other cells $c'$ of the same cell type that have at most one covariate different. Let $s_i$ be the covariate that differs in the pair $(c, c')$. Then, during training, we predict counterfactual gene expression values associated with $s_i$ (one $s_i$ at a time) and penalize them in the loss function using mean square error from the counterfactual pair.

#### 2 DISENTANGLED EXPERTS FOR COVARIATE COUNTERFACTUALS

##### 2.1 CELLDISECT ARCHITECTURE

Our approach to implementing abduction, action, and prediction is inspired by Foster et al. (2022), which streamlines it by imposing an orthogonality constraint between latent variables and observed covariates. However, their constraint needs to be generalized to meet the demands of our multi-covariate model. Our proposed generalization is the constraint $\forall i \neq j : \ z_i \perp\!\!\!\perp s_j$, which demands that different representations are aligned with only one counterfactual at a time.

The CellDISECT architecture and model has five main components : (1) encoders, (2) decoders, (3) the covariate classifiers, (4) the covariate adversarial classifiers, and (5) the semi-autoenciding of the counterfactuals.

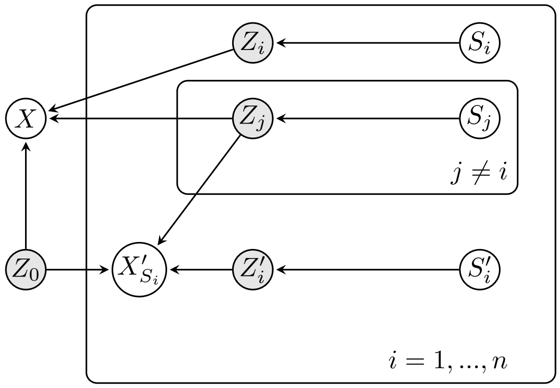


Figure 3: Many worlds graph for counterfactual prediction. There are $n$ worlds, each of which is associated with a single intervention.

#### 2.2 ENCODERS

We employ $n + 1$ encoders, parameterised by $\phi_0, \phi_1, \ldots, \phi_n$, to independently infer $n + 1$ latent vectors $z_0, z_1, \ldots, z_n$. Each encoder receives the same input: the gene expression vector $X$ and the complete covariate vector $s$.

Additionally, we use $n$ encoders, parameterised by $\psi_1, \ldots, \psi_n$, to infer $n$ prior distributions for the latent variables. The input to the encoder $\psi_k$ consists solely of $s_k$.

For the latent variable $z_k$, we apply amortised inference Kingma et al. (2019); Lopez et al. (2018) as follows:

$$q_{\phi_k}(z_k \mid x, s) \sim \text{Normal}\left(g_\mu(x), \text{diag}(g_\sigma^2(x))\right),$$

where Normal $(\mu, \sigma)$ is a Normal distirbution of mean $\mu$ and variance $\sigma$.

In line with Proposition 1 (proof found in Appendix Note X), we infer a prior distribution for $z_k$ as:

$$q_{\psi_k}(z_k \mid s_k) \sim \text{Normal}\left(g_\mu(s_k), \text{diag}(g_\sigma^2(s_k))\right).$$

#### 2.3 DECODERS

We implement $n + 1$ decoders, parameterised by $\theta_0, \theta_1, \ldots, \theta_n$, to generate the gene expression vector $x$ within each decoder independently. The input for decoder 0 is $(z_0; s)$, while for the $k$-th decoder (where $k \neq 0$), the input is $(z_k, s_{-k})$, where $s_{-k}$ refers to all attributes except $s_k$.

The rationale behind this setup is to ensure that $z_k$ encapsulates all the information in $s_k$ but none from $s_{-k}$, for $k \neq 0$ (see next section). This architecture enables CellDISECT to learn disentangled representations. Moreover, $z_0$ is designed to exclude any information about $s$, instead learning cell-specific background variation due to unobserved covariates.

Using decoder $\theta_k$, we predict the expression of gene $g$ (denoted $x_g$) using a Zero Inflated Negative Binomial (ZINB) distribution, or alternatively a Negative Binomial (NB) distribution, which has been shown to appropriately model gene expression data Lopez et al. (2018). The gene expression for gene $g$ in cell $j$ is given by:

$$x_g \sim \text{NegativeBinomial}\left(l f^g(z), d_g\right),$$

where $l$ is the observed random variable representing the number of RNA molecules captured for that gene in the cell. $f$ is a function that maps the latent space to the simplex of the gene expression space, and $d_g$ is the dispersion parameter of the negative binomial distribution. The ZINB distribution adds a mixture coefficient representing the weight of the point mass. By default, we use the ZINB distribution in the generative process, as it is well-suited for count data characterised by a large proportion of zeros, as is typical in gene expression data Lopez et al. (2018).

##### 2.4 COVARIATE CLASSIFIERS AND ADVERSARIAL CLASSIFIERS

We aim for the following properties in the latent space:

1. High $MI(z_i, s_i)$ for all $i \neq 0$ (i.e., high predictability of $s_i$ given $z_i$).
2. Low $MI(z_{-i}, s_i)$ for all $i \neq 0$ (i.e., low predictability of $s_i$ given $z_{-i}$).

Here, $MI$ refers to mutual information, defined as $MI(X,Y) = \sum_{x \in X} \sum_{y \in Y} P(x,y) \log\left(\frac{P(x,y)}{P(x)P(y)}\right)$.

To achieve property (1), for each $i \neq 0$, we employ a classifier to predict $s_i$ given $z_i$, and penalise the loss function with a cross-entropy loss $CE(\text{classifier}(s_i \mid z_i), s_i)$. To achieve property (2), for each $i \neq 0$ and $j \neq i$, we use a classifier to predict $s_i$ given $z_j$, and penalise the loss function with cross-entropy loss $CE(\text{classifier}(s_i \mid z_j), s_i)$.

All classifiers are fully connected networks with one hidden layer of size 128 by default, and they predict the one-hot encoding of the given attribute.

##### 2.5 LOSS FUNCTION

As explained in Appendix A (see Corollary 1), based on the structure of our assumed Structural Causal Model we derive a lower bound on the sum of the (factual) posterior likelihood and counterfactual posterior likelihood, which translates into the following loss function,

$$\begin{aligned}\mathcal{L}(\theta,\phi) = &-\alpha_1 \frac{1}{C+1} \sum_{i=0}^{C} \mathbb{E}_{q_{\phi_i}(z_i|x,S)} \log p_{\theta_i}(x|z_i, s_{-i}) - \alpha_2 \frac{1}{n_{cf}} \sum_{i=1}^{n_{cf}} \log \hat{p}(x'|x, s', s) \\ &+ \alpha_3 \frac{1}{C} \sum_{i=1}^{C} D_{\mathrm{KL}}\left(q_{\phi_i}(z_i|x,s) \,||\, q_{\psi_i}(z_i|s_i)\right) + \alpha_4 \sum_{i=1}^{C} CE(classifier(s_i|z_i), s_i) \\ &- \alpha_5 \mathcal{L}_{adv}(\theta,\phi)\end{aligned} \tag{28}$$

$$\mathcal{L}_{adv}(\theta,\phi) = \frac{1}{C^2} \sum_{i=1}^{C} \sum_{\substack{j=0 \\ j \neq i}}^{C} \mathrm{CE}\left(classifier_{ij}(s_i|z_j), s_i\right) \tag{29}$$

where $C$ is the number of observed covariates, $\hat{p}(x'|s')$ is the likelihood of the $x'$ in the counterfactual distribution derived from the method explained in Section 1.4, $n_{cf}$ is the number of counterfactual steps in the epoch, and $\alpha_1, \alpha_2, \alpha_3, \alpha_4, \alpha_5$ are heuristic hyperparameters which we explore in the ablation experiment.

#### 3 METRICS

##### 3.1 DISENTANGLEMENT METRICS

We divide disentanglement benchmarks into two scenarios. The first scenario has two covariates: a batch ID and a biological covariate. This is equivalent to a batch removal task in the field. The ideal method should remove the batch ID effect and preserve the biological covariate, which in this case is cell-type labels. We use standard metrics called scIB for each category as proposed by Luecken et al. (2022).

The second scenario extends the number of covariates beyond two, and only specific methods can handle such cases. We benchmark our method using the Mutual Information Gap [Chen et al. (2018); Higgins et al. (2016); Wu et al. (2023); Kumar et al. (2017); Kim & Mnih (2018)] due to its ability to generalize and be unbiased [Chen et al. (2018); Sepliarskaia et al. (2021)]. Specifically, we use two variations of MIG:

- maxMIG$(Z_1, ..., Z_n; S_1, .., S_n) = \frac{1}{n} \sum_{i=1}^{n} \frac{1}{H(S_i)} \max_{j \neq i} \Big[\mathrm{MI}(Z_i, S_i) - \mathrm{MI}(Z_i, S_j)\Big]$

- $\text{concatMIG}(Z_1, ..., Z_n; S_1, .., S_n) = \frac{1}{n}\sum_{i=1}^{n}\frac{1}{H(S_i)}\Big[\text{MI}(Z_i, S_i) - \text{MI}(Z_{-i}, S_i)\Big]$

It's worth noting that, by its very definition, catMIG can assume negative values. For example, when all the non-age latent spaces together are more informative about age than the age latent space.

#### REFERENCES


Ricky T. Q. Chen, Xuechen Li, Roger B Grosse, and David K Duvenaud. Isolating sources of disentanglement in variational autoencoders. In S. Bengio, H. Wallach, H. Larochelle, K. Grauman, N. Cesa-Bianchi, and R. Garnett (eds.), *Advances in Neural Information Processing Systems*, volume 31. Curran Associates, Inc., 2018. URL `https://proceedings.neurips.cc/paper_files/paper/2018/file/1ee3dfcd8a0645a25a35977997223d22-Paper.pdf`.

Adam Foster, Árpi Vezér, Craig A Glastonbury, Páidí Creed, Samer Abujudeh, and Aaron Sim. Contrastive mixture of posteriors for counterfactual inference, data integration and fairness. In *International Conference on Machine Learning*, pp. 6578–6621. PMLR, 2022.

C. W. J. Granger. Investigating causal relations by econometric models and cross-spectral methods. *Econometrica*, 37(3):424–438, 1969. ISSN 00129682, 14680262. URL `http://www.jstor.org/stable/1912791`.

Irina Higgins, Loic Matthey, Arka Pal, Christopher Burgess, Xavier Glorot, Matthew Botvinick, Shakir Mohamed, and Alexander Lerchner. beta-vae: Learning basic visual concepts with a constrained variational framework. In *International conference on learning representations*, 2016.

Ilyes Khemakhem, Diederik P. Kingma, Ricardo Pio Monti, and Aapo Hyvärinen. Variational autoencoders and nonlinear ica: A unifying framework, 2020. URL `https://arxiv.org/abs/1907.04809`.

Hyunjik Kim and Andriy Mnih. Disentangling by factorising. In Jennifer Dy and Andreas Krause (eds.), *Proceedings of the 35th International Conference on Machine Learning*, volume 80 of *Proceedings of Machine Learning Research*, pp. 2649–2658. PMLR, 10–15 Jul 2018. URL `https://proceedings.mlr.press/v80/kim18b.html`.

Diederik P Kingma, Max Welling, et al. An introduction to variational autoencoders. *Foundations and Trends® in Machine Learning*, 12(4):307–392, 2019.

Abhishek Kumar, Prasanna Sattigeri, and Avinash Balakrishnan. Variational inference of disentangled latent concepts from unlabeled observations. *arXiv preprint arXiv:1711.00848*, 2017.

Romain Lopez, Jeffrey Regier, Michael B Cole, Michael I Jordan, and Nir Yosef. Deep generative modeling for single-cell transcriptomics. *Nature methods*, 15(12):1053–1058, 2018.

Malte D Luecken, Maren Büttner, Kridsadakorn Chaichoompu, Anna Danese, Marta Interlandi, Michaela F Müller, Daniel C Strobl, Luke Zappia, Martin Dugas, Maria Colomé-Tatché, et al. Benchmarking atlas-level data integration in single-cell genomics. *Nature methods*, 19(1):41–50, 2022.

Judea Pearl. *Causality*. Cambridge university press, 2009.

Anna Sepliarskaia, Julia Kiseleva, and Maarten de Rijke. How to not measure disentanglement, 2021.

Yulun Wu, Layne C Price, Zichen Wang, Vassilis N Ioannidis, Robert A Barton, and George Karypis. Variational causal inference. *arXiv preprint arXiv:2209.05935*, 2022.

Yulun Wu, Layne C. Price, Zichen Wang, Vassilis N. Ioannidis, Robert A. Barton, and George Karypis. Variational causal inference, 2023.
